# Injury-induced nuclear export of RNA-binding proteins drives mRNA stabilization and translation to promote dendrite regeneration

**DOI:** 10.1101/2025.11.18.688889

**Authors:** Zhongwei Qu, Dong Yan

## Abstract

Dendrite regeneration is critical for restoring neuronal connectivity after injury, yet the underlying molecular mechanisms remain poorly understood. Using *C. elegans* as a model and through a forward genetic screen, we identified the conserved insulin degrading enzyme *idr-1* and the RNA-binding protein *rbm-42* as key regulators of dendrite regeneration, where *idr-1* functions upstream of *rbm-42*. We further show that *ced-7*, one of the core components of the phagocytosis pathway, acting down stream of *rbm-42*, while other components of this pathway don’t play significant roles in dendrite regeneration. In addition, we demonstrate that upon injury IDR-1 can promote RBM-42 nuclear export following injury, enabling its dendritic localization. RBM-42, in turn, promotes the translation of *ced-7* and facilitates microtubule assembly. In conclusion, our findings define a novel conserved signaling cascade coupling injury-induced nuclear export of RNA binding proteins to local regulation and dendrite regeneration, providing new mechanistic insight into neuronal repair.

## Introduction

Neurons depend on elaborate dendritic arbors to receive, integrate, and process synaptic inputs. Dendritic damage is a hallmark of many neurological disorders, including traumatic brain injury (*1*), stroke (*2*), epilepsy (*3*), and is closely associated with cognitive and behavioral deficits (*4*). In contrast to axonal regeneration, which has been studied extensively, the regenerative capacity of dendrites of different types of neurons and the underlying molecular mechanisms are largely unknown. Studies across different model organisms, including *C. elegans* (*5*), Drosophila (*6–9*), and mammals (*10*), have shown that dendrites can regrow and reconnect after injury, and dendrite regeneration use distinct mechanisms than axon reaeration (*9, 11, 12*). While a few genes have been identified contributing to dendrite regrowth upon injury, yet the underlying mechanisms that restore both structural integrity and functional output remain largely uncovered. Therefore, understanding dendrite regeneration at the molecular level will provide critical insight into neuronal resilience and repair.

RNA-binding proteins (RBPs) serve as central regulators of RNA processing, including splicing (*13, 14*), transportation (*15*), translation (*16–18*) and decay (*19*). In neurons, RBPs are particularly critical because of the polarized morphology of axons and dendrites, which requires spatially restricted and temporally precise protein synthesis. Local regulation of mRNAs guided by RBPs has been implicated in axon pathfinding (*20*), dendritic spine morphogenesis (*21*), and synaptic plasticity (*22*), ensuring that remodeling occurs in response to extracellular cues. Beyond developmental contexts, RBPs can shape axon regeneration by modulating mRNA stability and local translation (*23, 24*). Many transcripts encoding cytoskeletal regulators, membrane-trafficking proteins, and signaling molecules are selectively stabilized or translated near dendritic or axonal sites, enabling rapid remodeling in response to environmental signals (*25*). RBPs that recognize ARE or specific 3′-UTR motifs can determine whether an mRNA is degraded or locally translated, thereby influencing growth cone navigation (*26*), dendritic plasticity (*27*), and synaptic strength (*28*). Despite these advances, whether and how RBPs contribute to dendrite regeneration upon injury remains unknown.

The phagocytosis pathway was first identified in the context of apoptotic cell clearance and has since been shown to play essential roles in diverse biological processes (*29*). In the nervous system, phagocytic signaling maintains neuronal health by eliminating apoptotic cells and cellular debris (*30*), facilitates synaptic pruning (*31, 32*) and remodeling (*33, 34*), and provides protection against neurodegeneration (*35*). In addition, components of this pathway contribute to axonal fusion and repair after injury by mediating “find-me” and “eat-me” signals (*36*). Despite these established functions, it remains unclear whether and how phagocytosis-related signaling contributes to dendrite regeneration.

Here, using the mechanosensory PVD neurons of *C. elegans* as a model, we dissect the molecular mechanisms underlying dendrite regeneration. Through forward genetic screening and mechanistic analysis, we identify a conserved signaling axis composed of *idr-1,* the insulin degrading enzyme, *rbm-42,* an mRNA binding protein, and *ced-7,* one of the core components of the phagocytosis pathway, in regulating dendrite regeneration. We show that IDR-1 promotes injury-induced nuclear export of RBM-42, enabling its dendritic localization and post-transcriptional regulation of CED-7 via ARE in its 3′-UTR. Genetic analysis places these components in a linear pathway that accelerates dendritic outgrowth and stabilizes microtubules during regeneration. These findings reveal a previously unrecognized mechanism linking RNA regulation with microtubules dynamic to promote dendrite repair, providing new conceptual insight into the molecular basis of neuronal regeneration.

## Results

### *idr-1* is required for dendrite regeneration of PVD neurons in *C. elegans*

PVD neurons in *C. elegans* are mechanosensory neurons that share morphological features with nociceptors in mammals, utilize common gene sets for differentiation and function, and express ion channel components conserved across species (*37*) (Fig. 1A). These neurons have been validated as a reliable model for studying dendrite regeneration (*5*). To mimic acute dendritic injury, we performed dendrotomy on the primary dendrite of PVD neurons in 1-day-old adult animals using UV laser. Consistent with previous reports, regrowth typically occurs via outgrowth from the proximal stump followed by fusion with the distal fragment, which represents the predominant mode of recovery (Fig. 1A). Quantification revealed more than 60% of animals exhibited successful fusion at 24 hours post-dendrotomy, and this proportion progressively increased to >90% by 96 hours in control animals (Fig. 1B). To assess whether morphological reconnection restores function, we tested harsh-touch responses, a behavioral readout of PVD mechanosensory activity. To exclude contributions from gentle-touch receptor neurons, we conducted these assays in *mec-4* mutants, which lack gentle-touch sensation but retain harsh-touch responses mediated primarily by PVD neurons. We first excluded the potential effects of *mec-4* mutation on dendrite regeneration by examining the regenerative ability of loss of function *(lf)* in *mec-4* and found that *mec-4(lf)* did not affect dendrite regeneration (Fig. S1A). Next, we examined animals with and without dendritic fusion 24 hours after dendrotomy and found that animals whose dendrites successfully fused responded robustly to harsh touch, whereas those without dendritic fusion showed impaired responses (Fig. S1B). These findings demonstrate that dendritic reconnection in PVD neurons is both morphological and functional.

**Figure 1.**
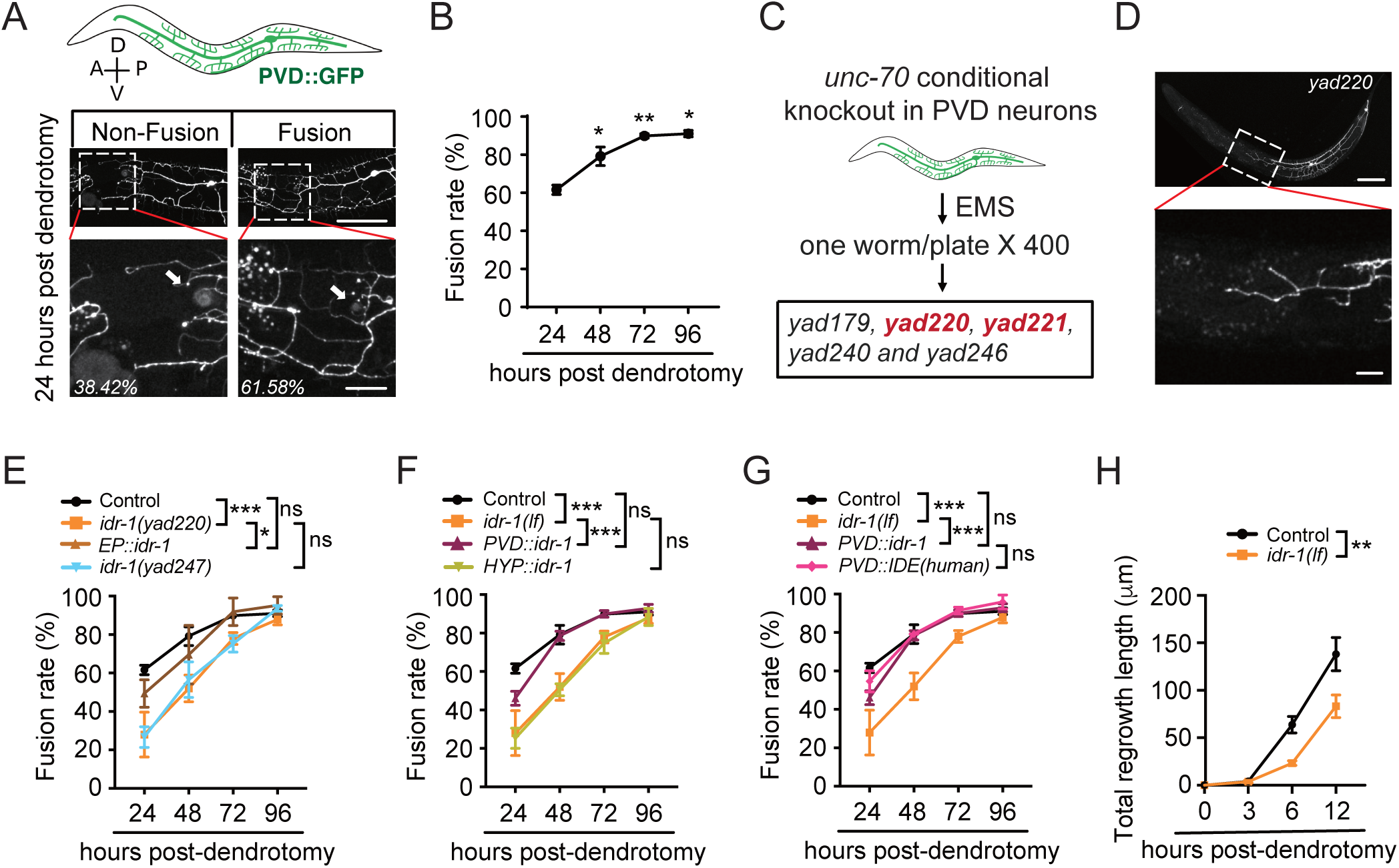
Use of *C. elegans* PVD neurons as a model to study dendrite regeneration. (A) Representative images of PVD neurons labeled with GFP. A: anterior; P: posterior; D: dorsal; V: ventral. Examples show non-fusion and fusion between the proximal and distal dendritic segments of PVD neurons upon UV-laser dendrotomy of control animals. White arrows indicate the injury sites. Scale bar, 100 μm (top) and 20 μm (enlarged views of boxed regions). (B) Quantification of the dendritic fusion rates in control animals at 24, 48, 72, and 96 hours post-dendrotomy. The dendritic fusion rate is defined as the percentage of animals in which regenerated dendrites fused with distal segments. *n*≥40. For all dendrotomy experiments, *n* in the figure legends throughout this paper indicates the number of animals per condition in each experiment, and statistics were derived from three independent biological experiments. (C) Schematic diagram illustrating the workflow of the forward genetic screen to identify mutants with defects in dendrite regeneration. A PVD–specific conditional knockout of *unc-70* strain was used as a genetic background to generate spontaneous breaks in the primary dendrites. After mutagenesis by ethyl methanesulfonate (EMS), 400 F1 animals (800 mutagenized haploid genomes) were screened, yielding five mutants with dendrite regeneration defects (*yad179*, *yad220*, *yad221*, *yad240* and *yad246*). (D) Representative image of the PVD neurons in *yad220* mutants, showing defective dendrite regeneration. Scale bar, 100 μm (top) and 20 μm (enlarged view of boxed region). (E) Quantification of dendritic fusion rates in control, *idr-1(yad220)*, *idr-1(yad247)*, and rescue transgenic strain with *idr-1(yad220)* background under its endogenous promotor (*EP::idr-1*). *idr-1(yad220)* is used as *idr-1(lf)* in the rest of this study unless noticed. *n*≥40. (F) Dendritic fusion rates of PVD neurons in control, *idr-1(lf)* and rescue transgenic strains. PVD- and hypodermis (HYP)-specific rescues were driven by *ser-2*(*3*) and *col-19* promotors, respectively. *n*≥32. (G) Quantification data show dendritic fusion rates in control, *idr-1(lf)* and rescue strains expressing *idr-1* or *IDE(human)* specifically in PVD neurons. *n*≥32. (H) Quantification data show the total regrowth lengths of the branches from the proximal stump after dendrotomy in control and *idr-1(lf)* animals. *n*≥15. Data represent mean ± standard deviation (SD). \**P* ≤ 0.05; \*\**P* ≤ 0.01; \*\*\**P* ≤ 0.001; ns, nonsignificant. Statistical significance was determined by one-way ANOVA followed by Tukey’s post hoc test in Fig. 1A and by two-way ANOVA followed by Tukey’s post hoc test in Fig. 1E-1H.

To identify molecules required for dendrite regeneration, we conducted an unbiased forward genetic screen. Given that loss of function in *unc-70*/β-spectrin induces spontaneous axonal breaks (*38*), we tested whether it also triggers dendritic breaks in PVD neurons. We generated a strain with conditional PVD-specific knockout of *unc-70* using the FLP/FRT recombination system and observed spontaneous breaks in the primary dendrite (Fig. S2A and S2B). Using this strain, we carried out a forward genetic screen and examined ∼800 mutagenized haploid genomes and isolated five mutants with defect in dendrite regeneration (Fig. 1C).

We further studied one of these mutants, *yad220*, which exhibited defects on dendrite regeneration (Fig. 1D). This phenotype was first confirmed by UV-laser induced dendrotomy in animals that removing the FLP/FRT recombination system and other genetic background through backcrossing against wild type animals. As shown in Fig. 1E, *yad220* animals displayed significantly reduced fusion rates at 24, 48, and 72 hours compared with controls (Fig. 1E). Whole-genome sequencing and mapping identified a single-nucleotide polymorphism (SNP) in a previously uncharacterized gene, *Y70C5C.1*, which we named it as *inducing dendrite regeneration-1 (idr-1)* (Fig. S3A). Expression of *idr-1* under its endogenous promoter (EP) rescued the defects in dendrite regeneration of *yad220* animals (Fig. 1E). To further validate the role of *idr-1*, we generated a null allele of *idr-1*, *yad247*, in which a frameshift mutation causes a 4-base-pair deletion and the premature stop codon (Fig. S3A). Our results showed that *yad247* phenocopied the regeneration defect caused by *yad220* (Fig. 1E), further supporting the function of *idr-1* in dendrite regeneration. Expression of *idr-1* specifically in PVD neurons, but not in hypodermal cells (HYP), restored regeneration, demonstrating that *idr-1* acts cell-autonomously in regulating dendrite regeneration (Fig. 1F).

We next investigated whether the function of *idr-1* is evolutionarily conserved by examining the rescue ability of its mammalian homolog, insulin-degrading enzyme (IDE) (Fig. S3A). The results showed that specific expression of human IDE in PVD neurons rescued the dendrite regeneration defects of *idr-1(lf)* mutants to a similar level as that in *C. elegans idr-1* transgenes, supporting the conserved role of the *idr-1* in dendrite regeneration across species (Fig. 1G). The classical function of IDE in substrate degradation has been attributed to its peptidase activity, which depends on several residues critical for catalysis (*39*). To test whether the enzymatic activity underlies *idr-1* function in dendrite regeneration, we mutated two conserved residues E73Q and Y460A in *idr-1* that are essential for its peptidase activity and expressed them in PVD neurons (Fig. S3A). Surprisingly, these transgenes retained full rescue ability (Fig. S3B and S3C), indicating that the peptidase activity is dispensable for *idr-1*-mediated dendrite regeneration. To further explore how *idr-1* promotes regeneration, we qualified dendritic outgrowth kinetics after injury. Regrowth from proximal stumps in *idr-1* mutants was significantly shorter than that in controls within the first 12 hours post-dendrotomy (Fig. 1H), suggesting that *idr-1* promotes dendrites extension during the early phase of regeneration.

### Loss of *rbm-42* suppressed dendrite regeneration of PVD neurons in *C. elegans*

We next focused on another mutant isolated in our generic screen, *yad221*. After backcross to remove all genetic backgrounds, we confirmed that *yad221* animals exhibited significantly reduced fusion rates compared with controls at 24, 48, and 72 hours post-dendrotomy (Fig. 2A and 2B). Whole-genome sequencing and genetic mapping identified a SNP in the gene *rbm-42* (Fig. S4A), the homolog of mammalian RNA-binding protein RBM42. Expression of *rbm-42* under its endogenous promoter not only rescued the regeneration defect of *rbm-42(lf)* animals but also generated a gain-of-function (*gf*) effect, enhancing dendrite regeneration beyond control levels (Fig. 2B). As the null alleles of *rbm-42* cause sterility and larval arrest, we first confirmed *yad221* is a loss-of-function allele by knocking down *rbm-42* specifically in PVD neurons through RNAi, which phenocopied the dendrite regeneration defects observed in *yad221* animals (Fig. 2C). Furthermore, expression of *rbm-42* specifically in PVD neurons rescued the dendrite regeneration defects in *rbm-42(lf)* animals, whereas expression in hypodermal cells had no effect, underscoring the cell-autonomous role of *rbm-42* in dendrite regeneration (Fig. 2D). Additionally, expression of the human RBM42 rescued regeneration defects in *rbm-42(lf)* mutants, demonstrating functional conservation between *rbm-42* and its mammalian homolog (Fig. 2E). To gain the insight in how *rbm-42* promotes dendrite regeneration, we quantified dendritic outgrowth after injury. Similar to that in *idr-1(lf)* mutants, *rbm-42(lf)* animals displayed significantly shorter regrowth lengths from the proximal stump when compared with controls in the first 12 hours post-dendrotomy (Fig. 2F), suggesting that *rbm-42* accelerates dendritic branch extension during the early phase of regeneration.

**Figure 2.**
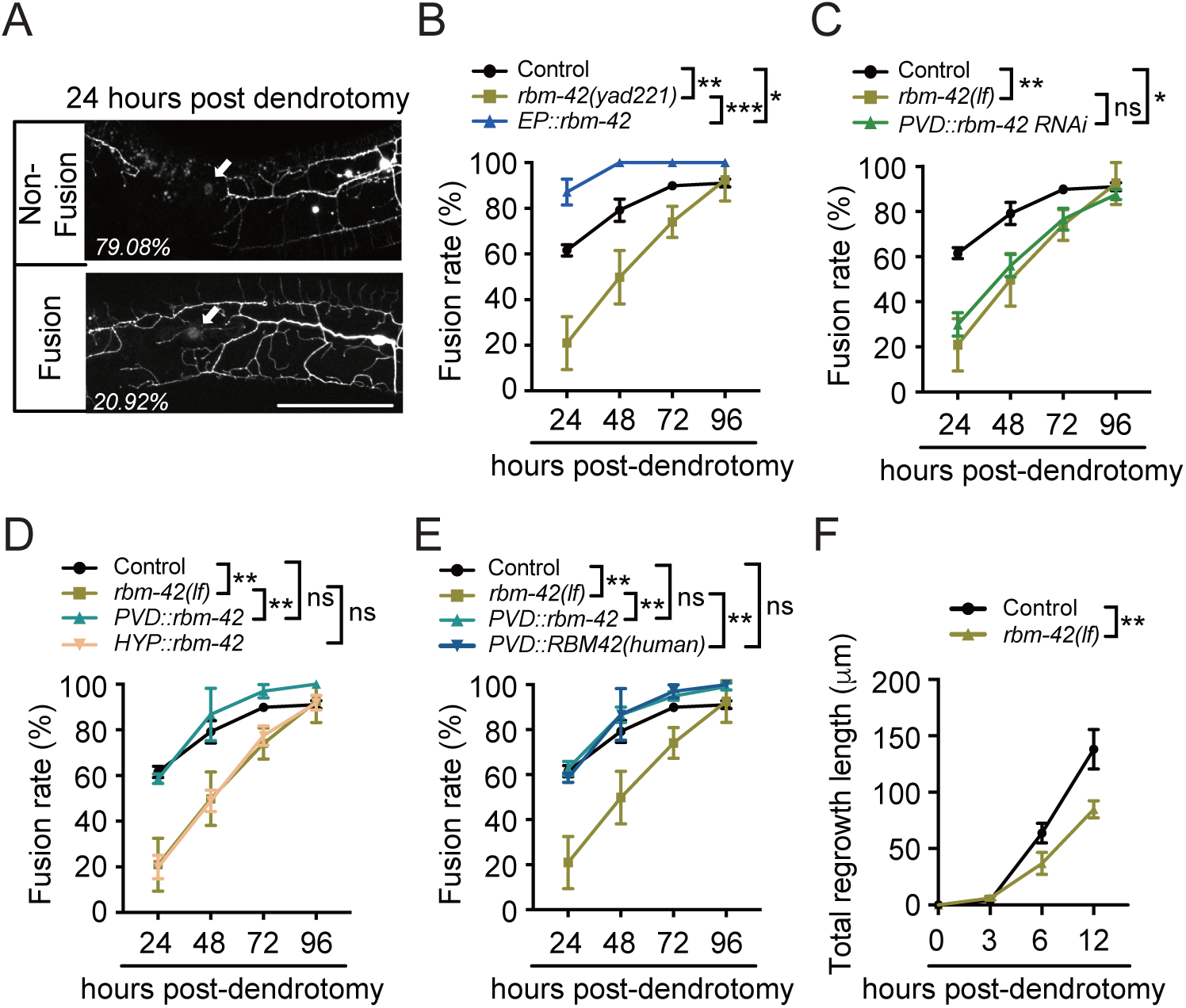
*rbm-42* is required for dendrite regeneration. (A) Representative images of PVD neurons in *rbm-42(yad221)* animals showing non-fusion and fusion phenotypes 24 hours post-dendrotomy. Scale bar, 100 μm. (B) Dendritic fusion rates in control, *rbm-42(yad221)*, and rescue strain expressing *rbm-42* under its endogenous promoter (*EP::rbm-42*). *rbm-42(yad221)* is used as *rbm-42(lf)* in the rest of this study unless noticed. Since *EP::rbm-42* causes a gain-of-function (*gf*) effect, it is referred to as *EP::rbm-42(gf)* throughout this study unless otherwise indicated*. n*≥28. (C) Quantification data show dendritic fusion rate in control, *rbm-42(lf)* and PVD-specific *rbm-42* knockdown animals. *n*≥28. (D) Quantification of dendritic fusion rates in control, *rbm-42(lf)* and rescue strains expressing *rbm-42* specifically in PVD neurons and hypodermal cells. *n*≥28. (E) Dendritic fusion rates in control, *rbm-42(lf)* and rescue strains expressing *rbm-42* or *RBM42(human)* specifically in PVD neurons. *n*≥28. (F) Quantification of total growth lengths of the branches from the proximal stump after dendrotomy in control and *rbm-42(lf)* animals. *n*≥15. Data represent mean ± SD from three independent experiments. \**P* ≤ 0.05; \*\**P* ≤ 0.01; \*\*\**P* ≤ 0.001; ns, nonsignificant. Statistical significance was determined by two-way ANOVA followed by Tukey’s post hoc test.

### *ced-7* played a key role in dendrite regeneration

In addition to performing the forward genetic screen, we also carried out a candidates-based screen, focusing on molecules previously implicated in homotypic adhesion and axonal fusion, including *ced-1*, *ced-7*, *ced-10* and *ina-1* (Fig. S5A). These genes are well known for their key roles in phagocytosis and in providing “fuse-me” signals during axonal repair (*40–43*), and INA-1 has been reported to function upstream of CED-10 to regulate axon regeneration (*44*). To test whether these factors also contribute to dendrite regeneration, we examined mutant strains using UV-laser dendrotomy. Dendritic fusion was almost completely abolished in *ced-7(lf)* animals (Fig. 3A and Fig. S5B), demonstrating that *ced-7* is essential for dendrite regeneration. In contrast, loss-of-function mutations in *ced-1*, *ced-10,* or *ina-1* had no detectable effect (Fig. S5C–E), suggesting that *ced-7* promotes regeneration and this process occurs independently of its canonical engulfment role. Because this finding contrasts with a previous report that *ced-7* is dispensable for dendrite repair (*45*), we tested a second null allele of *ced-7* and observed the same phenotype (Fig. 3A), confirming the essential role of *ced-7* in dendrite regeneration. In addition, expression of *ced-7* specifically in PVD neurons, but not in hypodermal cells, rescued the regeneration defect in two *ced-7* null alleles (Fig. 3B, S5F), indicating that *ced-7* functions in a cell-autonomous manner. However, because distal dendrite fragments degenerate rapidly after severing, rescue by the *ced-7* transgene did not fully restore fusion rates. To further validate the cell-autonomous role of *ced-7* in dendrite regeneration, we specifically knocked down *ced-7* in PVD neurons and found similar reduction of dendrite regeneration (Fig. S5G). We next investigated how *ced-7* contributes to dendrite regeneration. After measuring dendritic regrowth kinetics, we found that *ced-7(lf)* animals exhibited significantly shorter proximal regrowth compared with controls (Fig. 3C), indicating that *ced-7* promotes dendrite outgrowth upon injury.

**Figure 3.**
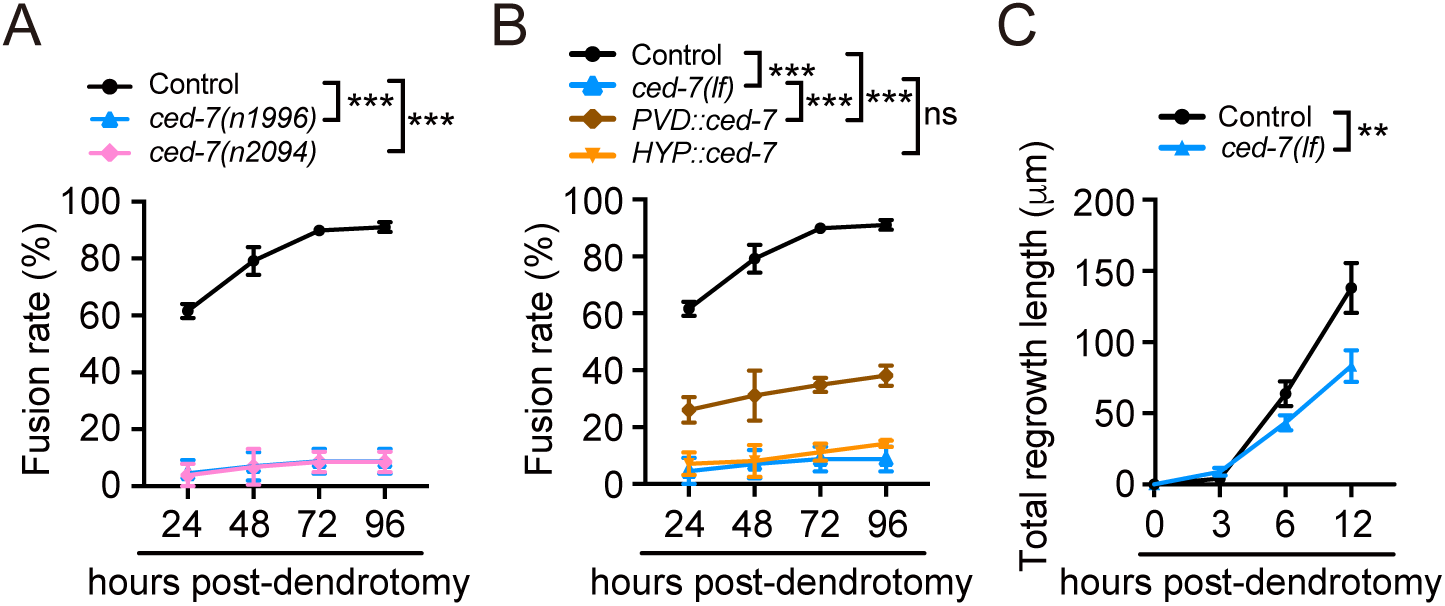
Loss of function of *ced-7* suppresses dendrite regeneration. (A) Quantification data show dendritic fusion rates in control, *ced-7(n1996)* and *ced-7(n2094)* animals. *ced-7(n1996)* is used as *ced-7(lf)* in subsequent experiments unless otherwise noticed. *n*≥24. (B) Dendritic fusion rates in control, *ced-7(lf)* and rescue strains expressing *ced-7* in PVD neurons or hypodermal cells, respectively. *n*≥24. (C) Quantification data show the total regrowth lengths of the branches from the proximal stump after dendrotomy in control and *ced-7(lf)* animals. *n*≥16. Data represent mean ± SD. \*\**P* ≤ 0.01; \*\*\**P* ≤ 0.001; ns, nonsignificant. Statistical significance was determined by two-way ANOVA followed by Tukey’s post hoc test.

### *idr-1*, *rbm-42* and *ced-7* functions in a same genetic pathway to regulate dendrite regeneration

Having established that *idr-1*, *rbm-42*, and *ced-7* each play critical roles in dendrite regeneration, we next investigated their genetic relationships to determine whether they function in a linear pathway. Double mutants of *idr-1(yad247)* and *rbm-42(lf)* did not display enhanced defects in dendritic fusion and regrowth length compared with *idr-1(yad247)* alone (Fig. 4A and 4B), suggesting that *idr-1* and *rbm-42* act in the same genetic pathway. As both *rbm-42* and *ced-7* are located on the same chromosome, we examined *rbm-42(lf)ced-7(lf)* double mutant phenotypes by introduced the *ced-7(yad306)* mutation into the *rbm-42(yad221)* animals using CRISPR. Since *yad306* caused one-base-pair deletion in *ced-7*, resulting in a frameshift and a premature stop codon (Fig. S5B), thus it likely represents a null allele. Our results show *rbm-42(yad221)ced-7(yad306)* double mutants did not exacerbate regeneration defects when compared with *ced-7(lf)* alone (Fig. 4C and 4D), indicating that *rbm-42* and *ced-7* function in the same genetic pathway.

**Figure 4.**
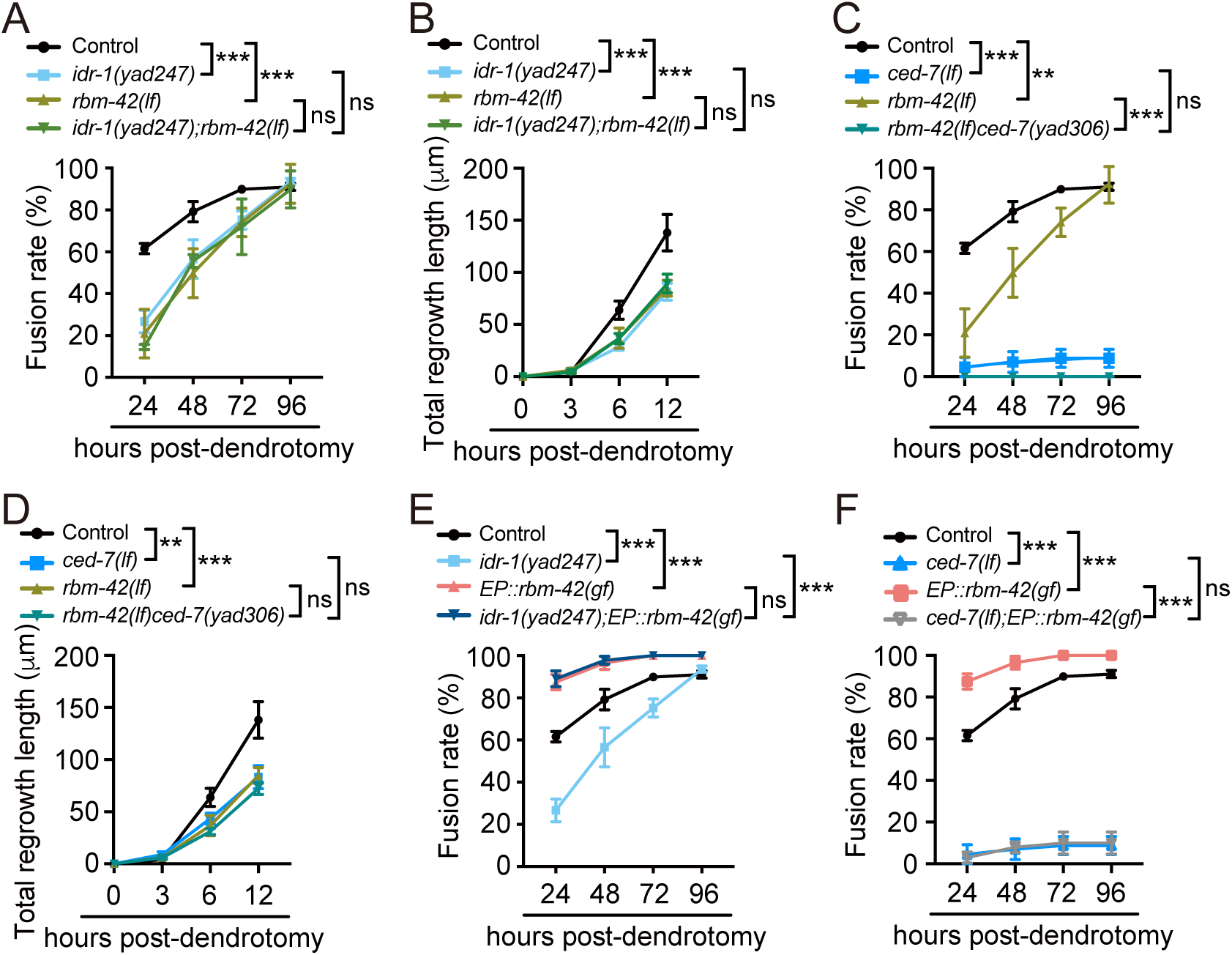
*idr-1, rbm-42* and *ced-7* functions in the same genetic pathway. (A) Quantification data show dendritic fusion rates in control, *idr-1(yad247), rbm-42(lf)* and *idr-1(yad247);rbm-42(lf)* double mutant animals. *n*≥28. (B) Quantification data show total regrowth lengths from the proximal stump after dendrotomy was measured in control, *idr-1(yad247), rbm-42(lf)* and *idr-1(yad247);rbm-42(lf)* double mutant animals. *n*≥15. (C) Dendritic fusion rates in control, *rbm-42(lf)*, *ced-7(lf)* and *rbm-42(lf)ced-7(yad306)* double mutant animals. *n*≥24. (D) Quantification of the total regrowth lengths of dendritic branches from the proximal stump after dendrotomy in control, *rbm-42(lf)*, *ced-7(n1996)* and *rbm-42(lf) ced-7(yad306)* double mutant animals. *n*≥15. (E) Dendritic fusion rates in control, *idr-1(yad247), EP::rbm-42(gf)* and *idr-1(yad247);EP::rbm-42(gf)* transgene animals. *n*≥28. (F) Quantification data show dendritic fusion rates in control, *ced-7(lf), EP::rbm-42(gf)* and *ced-7(lf);EP::rbm-42(gf)* transgene animals. *n*≥24. Data represent mean ± SD from three independent experiments. \*\**P* ≤ 0.01; \*\*\**P* ≤ 0.001; ns, nonsignificant. Statistical significance was determined by two-way ANOVA followed by Tukey’s post hoc test.

To determine the relationship among *idr-1, rbm-42*, and *ced-7*, we crossed *rbm-42(gf)* transgenes into *idr-1(lf)* and *ced-7(lf)* mutants, respectively. The results showed that *idr-1(lf)* didn’t suppress the regeneration-promoting effects of *rbm-42(gf)* transgenes (Fig. 4E), placing *rbm-42* downstream of *idr-1*. In contrast, *ced-7(lf)* completely suppressed the gain-of-function effect of *rbm-42(gf)* (Fig. 4F), positioning *ced-7* downstream of *rbm-42* in dendrite regeneration (Fig. S5H). In conclusion, we demonstrated that *idr-1*, *rbm-42* and *ced-7* function in the same genetic pathway, in which *idr-1* functions upstream of *rbm-42,* while *ced-7* acts downstream of *rbm-42*.

### The IDR-1-dependent nuclear export of RBM-42 is required for dendrite regeneration

Next, we examined the regulatory mechanisms between *rbm-42* and *idr-1*. We first examined how RMB-42 response to injury by expressing a functional reporter *rbm-42::gfp* in PVD neurons. Without injury, in control animals about 60% of RBM-42 proteins are localized into the nucleus, and rest of them are distributed in soma and dendrites (Fig. 5A and 5B). Following dendrotomy, RBM-42 translocated out of the nuclear within 3 hours and clustered at the proximal injury site (Fig. 5A-5C), while the overall expression of RMB-42 remains unchanged (Fig. 5D). These nuclear export and dendritic local clusters of RMB-42 required *idr-1*, as *idr-1(lf)* blocked those injury-induced phenotypes. The effect of *idr-1(lf)* is not due to alternation of RMB-42 expression, as the overall expression of RMB-42 is comparable in control and *idr-1(lf)* animals (Fig. 5A-5D). With these findings, we further explored the functional significance of this RBM-42 nuclear export in dendrite regeneration. Toward this goal, we added two repeated nuclear localization sequence (nls) of SV40 at the N-terminal of *rbm-42::gfp* and expressed it in PVD neurons (Fig. S4B). This modification caused RMB-42 to localize exclusively in the nuclear and abolishes the injury-dependent nuclear translocation of RMB-42 (Fig. 5E). We then examined the rescue ability of this transgene and found it failed to rescue *rmb-42(lf)* phenotypes (Fig. 5F), supporting the essential role of RBM-42 nuclear export in injury-triggered dendrite regeneration. Taking together, these results demonstrate that the critical role of RBM-42 nuclear export in dendrite regeneration and highlight the function of *idr-1* in regulating this injury-induced process.

**Figure 5.**
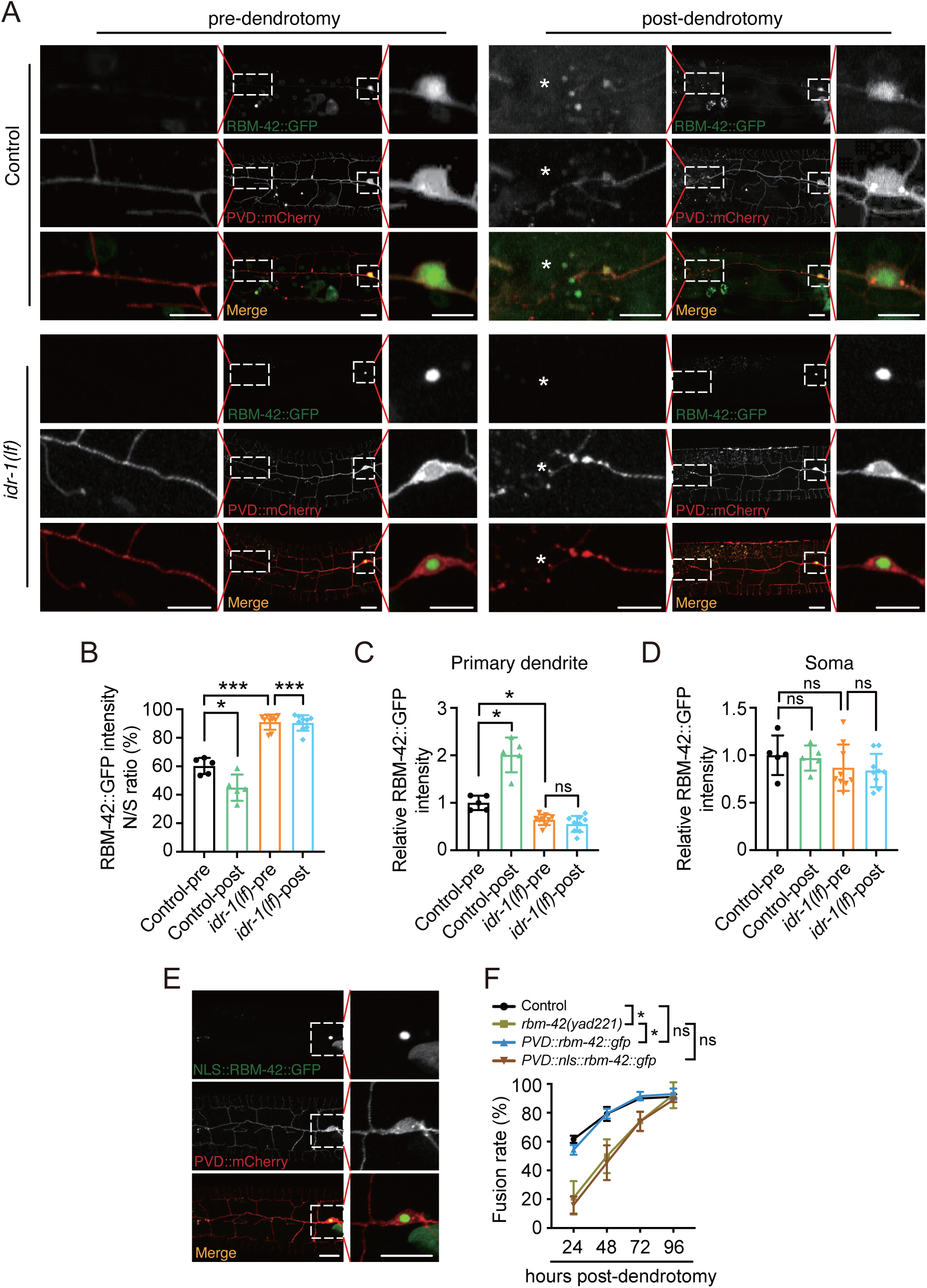
Injury-dependent RBM-42 nuclear export mediated by IDR-1 is required for dendrite regeneration. (A) Representative images illustrating the nuclear localization of RBM-42::GFP in control and *idr-1(lf)* animals pre- and 3 hours post-dendrotomy. Free mCherry was co-expressed with RBM-42::GFP in PVD neurons to label the neuronal morphology. Enlarged views show RBM-42::GFP distribution in the primary dendrite at the injury site (*) and in soma. Scale bar, 20 μm. (B) Quantification of the nuclear-to-soma (N/S) ratio of RBM-42::GFP intensity in control and *idr-1(lf)* animals pre- and post-dendrotomy. Nuclear RBM-42::GFP intensity was normalized to that in the soma. *n*=5. (C) Relative RBM-42::GFP intensity in the primary dendrite at injury sites (20 μm) in control and *idr-1(lf)* animals pre- and post-dendrotomy. RBM-42::GFP intensity was normalized to mCherry and further normalized to the mean RBM-42::GFP/mCherry ratio in control animals before dendrotomy. *n*=5. (D) Quantification of relative RBM-42::GFP intensity in the soma of control and *idr-1(lf)* animals pre- and 3 hours post-dendrotomy. RBM-42::GFP intensity was normalized to mCherry and further normalized to the mean RBM-42::GFP/mCherry ratio in control animals before dendrotomy. *n*=5. (E) Representative images showing nuclear localization of RBM-42::GFP fused with two tandem SV40 nuclear localization sequences (NLS). mCherry was co-expressed with NLS::RBM-42::GFP in PVD neurons to label neuronal morphology. Enlarged views highlight nuclear localization. Scale bar, 20 μm. (F) Dendritic fusion rates in control, *rbm-42(lf)* and rescue strains expressing *PVD::rbm-42::gfp* or *PVD::nls::rbm-42::gfp*, respectively. *n*≥28. Data represent mean ± SD from three independent experiments. \**P* ≤ 0.05; \*\*\**P* ≤ 0.001; ns, nonsignificant. Statistical significance was determined by one-way ANOVA followed by Tukey’s post hoc test in Fig.5B-5D and by two-way ANOVA followed by Tukey’s post hoc test in Fig.5F.

### RBM-42 regulates CED-7 expression

Next, we further explored how RBM-42 regulates CED-7 in dendrite regeneration. As RBM-42 is an RNA-binding protein known to regulate splicing and translation of its targets (*46, 47*), to test whether RBM-42 controls dendrite regeneration via RNA binding, we deleted the RNA-binding motif (ΔRBM) from *rbm-42* and examined its rescue ability (Fig. S4A). Our results showed that expression of *rbm-42* without the RNA-binding motif (RBM) failed to rescue the regeneration defects of *rbm-42(lf)* (Fig. 6A), supporting the essential role of RNA binding ability of RBM-42 in dendrite regeneration. Given that *ced-7* acts downstream of *rbm-42*, we next examined whether *ced-7* expression is regulated by RBM-42. Quantitative analysis revealed that *ced-7* mRNA levels were reduced in *idr-1(lf)* and *rbm-42(lf)* animals, whereas *rbm-42(gf)* increased *ced-7* expression (Fig. 6B), suggesting that the expression of *ced-7* is positively regulated by the *idr-1–rbm-42* pathway.

**Figure 6.**
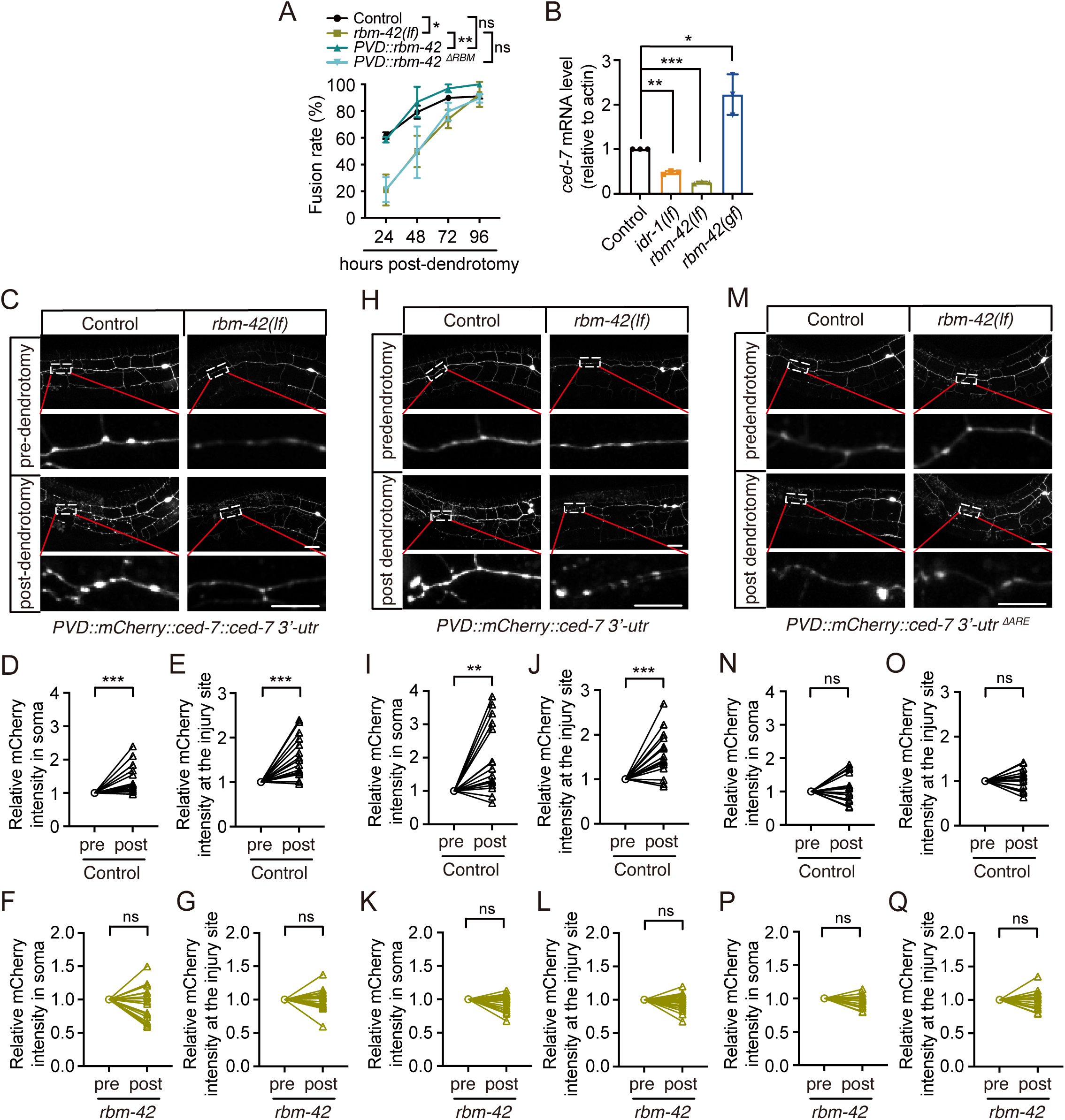
RBM-42 regulates the expression of CED-7. (A) Quantification of dendritic fusion rates in control, *rbm-42(lf)* and rescue strains expressing either full-length *rbm-42* (*PVD::rbm-42*) or *rbm-42* lacking its RNA-binding motif (*PVD::rbm-42* ^Δ*RBM*^). *n*≥28. (B) Relative *ced-7* mRNA levels in control, *idr-1(lf)*, *rbm-42(lf)*, and *rbm-42(gf)* animals measured by RT–qPCR. *n*=3. (**C**-**G**) Representative images (C) and quantification of *mCherry::ced-7::ced-7 3’-UTR* relative intensity in the soma and at the injury sites (20 μm) 6 hours post-dendrotomy in control (D: soma; E: injury site) and *rbm-42(lf)* (F: soma; G: injury site) animals. Post-dendrotomy (post) mCherry intensity in each group was normalized to the corresponding pre-dendrotomy (pre) value. The injury sites were boxed and enlarged. Scale bar, 20 μm. *n*≥17. (**H**-**L**) Representative images (H) and quantification of *mCherry::ced-7 3’-UTR* relative intensity in the soma and at the injury sites 6 hours post-dendrotomy in control (I: soma; J: injury site) and *rbm-42(lf)* (K: soma; L: injury site) animals. Post-dendrotomy (post) mCherry intensity in each group was normalized to the corresponding pre-dendrotomy (pre) value. The injury sites were boxed and enlarged. Scale bar, 20 μm. *n*≥17. (**M**-**Q**) Representative images (M) and quantification of *mCherry::ced-7 3’-UTR* ^Δ*ARE*^ relative intensity in the soma and at the injury sites 6 hours post-dendrotomy in control (N: soma; O: injury site) and *rbm-42(lf)* (P: soma; Q: injury site) animals. Post-dendrotomy (post) mCherry intensity in each group was normalized to the corresponding pre-dendrotomy (pre) value. The injury sites were boxed and enlarged. Scale bar, 20 μm. *n*≥17. Data represent mean ± SD from three independent experiments. \**P* ≤ 0.05; \*\**P* ≤ 0.01; \*\*\**P* ≤ 0.001; ns, nonsignificant. Statistical significance was determined by two-way ANOVA followed by Tukey’s post hoc test for Fig. 6A, by one-way ANOVA followed by Tukey’s post hoc test for Fig. 6B, by paired Student’s t-test for the rest statistical analysis.

To determine how RBM-42 regulates *ced-7* protein levels during regeneration, we expressed *ced-7* with its own 3’ UTR fused with mCherry in PVD neurons (Fig. 6C and Fig. S6A). Given the complex morphology of PVD neurons, we measured the CED-7 expression in soma and at the injury sites to assess alterations of its protein levels. In control animals, CED-7 expression increased both in soma and at the injury sites post-dendrotomy. (Fig. 6C-6E). In contrast, dendrotomy failed to induce CED-7 upregulation either in soma or at the injury sites in *rbm-42(lf)* animals (Fig. 6C, 6F and 6G). Since RMB-42 can regulate *ced-7* mRNA level, and it is known to regulate its target mRNA level through binding with the AU-rich element (ARE) in their 3′-UTRs to control RNA stability or translation (*47*), we next fused mCherry to the 3′-UTR of *ced-7* and expressed it in PVD neurons (Fig. S6A). Our results showed that dendrotomy increased mCherry intensity both in soma and at the injury sites in control animals (Fig. 6H -6J), whereas *rbm-42(lf)* abolished injury-induced mCherry intensity upregulation (Fig. 6K and 6L). These findings indicate that RBM-42 regulates *ced-7* via its 3′-UTR. As there is one ARE sequence in the 3′-UTR of *ced-7*, to probe the mechanism further, we deleted this ARE from the *ced-7* 3′-UTR (Fig. S6A and S6B) and found that deletion of the ARE prevented dendrotomy-induced increases of mCherry signal intensity either in soma or at the injury sites of control animals (Fig. 6M-6O). Moreover, ARE deletion didn’t affect the expression of mCherry in *rbm-42(lf)* animals post-dendrotomy (Fig. 6M, 6P and 6Q). Taken together, these results demonstrate that RBM-42 regulates *ced-7* expression through ARE-dependent interactions within the *ced-7* 3′-UTR.

### The *idr-1-rbm-42-ced-7* pathway is required for microtubule assembly during regeneration

As loss of function of all these three genes suppresses the growth of regenerating dendrites, a process that critically depends on the extension and stabilization of microtubules. Thus, we examined whether mutation of these genes affects microtubule assembly. To achieve this aim, we used a well-established marker for polymerized microtubes, EMTB::GFP, which fused the microtubule-binding domain of ensconsin (EMTB) with GFP (*48, 49*). In control animals, EMTB::GFP was evenly distributed along the primary dendrites without injury, but quickly accumulated at injury sites within 6 hours after dendrotomy (Fig. 7A), indicating increased microtubule stabilization at regenerating dendritic tips. As shown in our data, *idr-1(lf)*, *rbm-42(lf)*, and *ced-7(lf)* all reduced regrowth speed of proximal branches (Fig. 1H, 2F and 3C), we next tested whether these genes regulate microtubule stabilization. The results showed that *idr-1(lf)*, *rbm-42(lf)*, and *ced-7(lf)* all suppressed the accumulation of EMTB::GFP at dendrotomy sites post-dendrotomy (Fig. 7B), demonstrating that the *idr-1–rbm-42–ced-7* pathway regulates dendrite regeneration through promoting microtubule stabilization post-dendrotomy (Fig. S7A).

**Figure 7.**
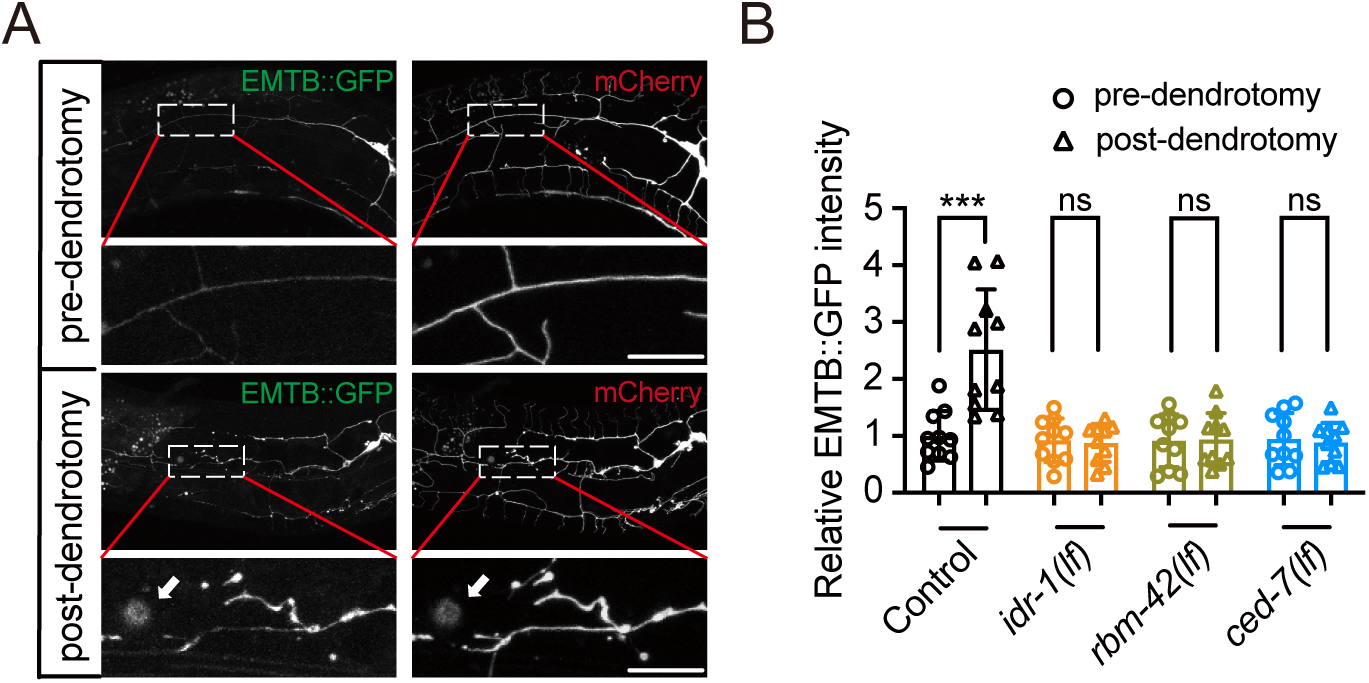
*idr-1-rbm-42-ced-7* pathway controls microtubule stabilization. (A) Representative images of PVD neurons labeled with a polymerized microtubule marker EMTB::GFP pre- and 6 hours post-dendrotomy. Free mCherry was co-expressed in PVD neurons to label the neuronal morphology. White arrow indicates the injury site. Enlarged views of the boxed regions show the EMTB::GFP enrichment at the injury site. Scale bar, 20 μm. (B) Quantification of EMTB::GFP intensity at injury sites (20 μm) in control, *idr-1(lf)*, *rbm-42(lf)* and *ced-7(lf)* animals pre-dendrotomy and 6 hours post-dendrotomy. Post-dendrotomy EMTB::GFP intensity in each group was normalized to the corresponding pre-dendrotomy value. *n*=10. Data represent mean ± SD. \*\*\**P* ≤ 0.001; ns, nonsignificant. Statistical significance was determined by paired Student’s t-test.

## Discussion

Dendritic injury is a prominent feature of many neurological disorders (*1–4*). However, the cellular and molecular basis of dendrite repair remains far less understood than axonal regeneration. Here, we identify a conserved pathway consisting of *idr-1*, *rbm-42*, and *ced-7* that promotes dendritic outgrowth and reconnection through regulating the microtubules assembly at the injury site in *C. elegans*. Our study identifies a new signal pathway mediating dendrite regeneration and expands the conceptual landscape of neuronal repair.

Our study also uncovers the conserved role of *idr-1* and its mammalian homolog IDE in mediating dendrite regeneration through an unexpected manner. While IDE has been primarily characterized as a peptidase that degrades insulin and amyloid peptides (*39, 50*), our genetic analysis shows that its peptidase activity is not required for dendrite regeneration. Instead, IDR-1 regulates RBM-42 nuclear export, placing *idr-1* upstream of RBP-mediated RNA regulation. This novel finding is supplement to previous work that emphasized IDE’s enzymatic function in metabolic regulation and amyloid clearance and instead suggests a non-canonical role for this conserved protein family in neuronal repair. Importantly, these results raise the possibility that IDE may contribute to dendritic repair in mammalian neurons, and in a novel fashion.

RBPs are well recognized for their roles in neuronal development, dendritic morphogenesis, and synaptic plasticity, largely through regulation of splicing, mRNA transport, and local translation (*20–22, 51*). However, their contribution to dendrite regeneration has not been explored. Our findings demonstrate that RBM-42 acts as a key post-transcriptional regulator of dendritic repair. In particular, dendrotomy triggers RBM-42 nuclear export and dendritic accumulation, a process dependent on *idr-1*. This injury-induced relocalization enables RBM-42 to regulate *ced-7* expression and thereby promote dendrite regeneration. These results extend the function of RBPs from previously well-defined developmental processes and axonal regeneration to injury-induced dendrites repair, highlighting RNA metabolism as a central node in neuronal resilience. Post-transcriptional control allows neurons to rapidly adjust protein synthesis in a spatially restricted manner (*52, 53*). Many RBPs are known to influence stability or local translation of transcripts encoding cytoskeletal or signaling proteins (*25*). Our data show that RBM-42 positively regulates *ced-7* expression via AU-rich elements in its 3′-UTR, thereby coupling mRNA stability with dendritic repair. Notably, while canonical RBPs often regulate growth cone guidance or synaptic remodeling, our findings discover a distinct injury-triggered program in which nuclear export and dendritic relocalization of an RBP directly promote regeneration. This mechanism by which RBM-42 exerts its critical role in dendrite repair greatly extends the current understanding of other RBPs previously studied in developmental contexts.

Our findings reveal that injury triggers nuclear export of RBM-42, enabling its dendritic localization and subsequent regulation of CED-7 expression. How RBM-42 promotes CED-7 upregulation at regenerating tips remains an open question. One possibility is that RBM-42 accumulates at the dendritic tip firstly followed by recruiting *ced-7* mRNA and facilitates its local translation, thereby ensuring rapid and spatially restricted production of CED-7 protein at injury sites. Alternatively, RBM-42 may bind *ced-7* mRNA in the soma and co-transport the ribonucleoprotein complex into dendrites, where translation is subsequently initiated. Both models are possible with our observation that RBM-42 requires its RNA-binding motif and the *ced-7* 3′-UTR ARE for mediating dendrite regeneration. Distinguishing between these scenarios will be critical for understanding whether RBM-42 primarily acts as a local translational enhancer or as an mRNA transport factor in dendrite regeneration.

Together, our findings define an *idr-1–rbm-42–ced-7* pathway that links RNA regulation and cytoskeletal dynamics to promote dendrite regeneration. This pathway provides a mechanistic framework for how injury-induced signals are transduced into local molecular programs that enable structural and functional repair. More broadly, these findings suggest that post-transcriptional regulation may represent general strategies employed by neurons to restore connectivity after damage.

## Acknowledgements

We thank all Yan lab members for comments on the manuscript and double-blind experiments. We are grateful to NBRP and CGC for providing us *C. elegans* strains, to Sunybiotech (Fujian, China) for generating FRT knock in strain. We appreciate Dr. Zheng Zhou (Baylor College of Medicine) for *ced-7* plasmid and DNASU (Arizona State University) for human IDE and RBM42 plasmids. We thank Dr. Bo Yuan (Institute of Medical Genetics and Genomics, Fudan University) for whole genomic sequencing data analysis. Yuanjun Yin, Yuzi Wang and Meghan Quinlan assisted the preparation of mutant strains and plasmids.

## Author Contributions

Z.Q. and D.Y. conceived the study and interpreted all the data. Z.Q. performed most of the experiments and data collection. D.Y. oversaw all experiments and data interpretation. Z.Q. and D.Y. wrote the manuscript.

## Data and materials availability

All data are available in the main text or the supplementary materials.

## Reference

1. X. Gao, J. Chen, Mild traumatic brain injury results in extensive neuronal degeneration in the cerebral cortex. J Neuropathol Exp Neurol 70, 183–191 (2011).

2. W. C. Risher, D. Ard, J. Yuan, S. A. Kirov, Recurrent spontaneous spreading depolarizations facilitate acute dendritic injury in the ischemic penumbra. J Neurosci 30, 9859–9868 (2010).

3. M. Isokawa, M. F. Levesque, Increased NMDA responses and dendritic degeneration in human epileptic hippocampal neurons in slices. Neurosci Lett 132, 212–216 (1991).

4. E. Falke et al., Subicular dendritic arborization in Alzheimer’s disease correlates with neurofibrillary tangle density. Am J Pathol 163, 1615–1621 (2003).

5. M. Oren-Suissa, T. Gattegno, V. Kravtsov, B. Podbilewicz, Extrinsic Repair of Injured Dendrites as a Paradigm for Regeneration by Fusion in Caenorhabditis elegans. Genetics 206, 215–230 (2017).

6. Y. Song et al., Regeneration of Drosophila sensory neuron axons and dendrites is regulated by the Akt pathway involving Pten and microRNA bantam. Genes Dev 26, 1612–1625 (2012).

7. L. DeVault et al., Dendrite regeneration of adult Drosophila sensory neurons diminishes with aging and is inhibited by epidermal-derived matrix metalloproteinase 2. Genes Dev 32, 402–414 (2018).

8. Y. Kitatani et al., Drosophila miR-87 promotes dendrite regeneration by targeting the transcriptional repressor Tramtrack69. PLoS Genet 16, e1008942 (2020).

9. M. C. Stone, R. M. Albertson, L. Chen, M. M. Rolls, Dendrite injury triggers DLK-independent regeneration. Cell Rep 6, 247–253 (2014).

10. J. Agostinone et al., Insulin signalling promotes dendrite and synapse regeneration and restores circuit function after axonal injury. Brain 141, 1963–1980 (2018).

11. K. Rao et al., Spastin, atlastin, and ER relocalization are involved in axon but not dendrite regeneration. Mol Biol Cell 27, 3245–3256 (2016).

12. D. M. R. Nye et al., The receptor tyrosine kinase Ror is required for dendrite regeneration in Drosophila neurons. PLoS Biol 18, e3000657 (2020).

13. J. Hang, R. Wan, C. Yan, Y. Shi, Structural basis of pre-mRNA splicing. Science 349, 1191–1198 (2015).

14. A. Ergun et al., Differential splicing across immune system lineages. Proc Natl Acad Sci U S A 110, 14324–14329 (2013).

15. E. Matoulkova, E. Michalova, B. Vojtesek, R. Hrstka, The role of the 3’ untranslated region in post-transcriptional regulation of protein expression in mammalian cells. RNA Biol 9, 563–576 (2012).

16. M. S. Lee, B. Kim, G. T. Oh, Y. J. Kim, OASL1 inhibits translation of the type I interferon-regulating transcription factor IRF7. Nat Immunol 14, 346–355 (2013).

17. W. J. Lin et al., Posttranscriptional control of type I interferon genes by KSRP in the innate immune response against viral infection. Mol Cell Biol 31, 3196–3207 (2011).

18. B. Herdy et al., The RNA-binding protein HuR/ELAVL1 regulates IFN-beta mRNA abundance and the type I IFN response. Eur J Immunol 45, 1500–1511 (2015).

19. D. R. Schoenberg, L. E. Maquat, Regulation of cytoplasmic mRNA decay. Nat Rev Genet 13, 246–259 (2012).

20. Y. Saito et al., NOVA2-mediated RNA regulation is required for axonal pathfinding during development. Elife 5, (2016).

21. M. L. Moeller, Y. Shi, L. F. Reichardt, I. M. Ethell, EphB receptors regulate dendritic spine morphogenesis through the recruitment/phosphorylation of focal adhesion kinase and RhoA activation. J Biol Chem 281, 1587–1598 (2006).

22. K. Si, Y. B. Choi, E. White-Grindley, A. Majumdar, E. R. Kandel, Aplysia CPEB can form prion-like multimers in sensory neurons that contribute to long-term facilitation. Cell 140, 421–435 (2010).

23. P. Patel et al., Intra-axonal translation of Khsrp mRNA slows axon regeneration by destabilizing localized mRNAs. Nucleic Acids Res 50, 5772–5792 (2022).

24. P. K. Sahoo et al., A Ca(2+)-Dependent Switch Activates Axonal Casein Kinase 2alpha Translation and Drives G3BP1 Granule Disassembly for Axon Regeneration. Curr Biol 30, 4882–4895 e4886 (2020).

25. Y. Fonkeu et al., How mRNA Localization and Protein Synthesis Sites Influence Dendritic Protein Distribution and Dynamics. Neuron 103, 1109–1122 e1107 (2019).

26. K. Y. Wu et al., Local translation of RhoA regulates growth cone collapse. Nature 436, 1020–1024 (2005).

27. D. M. Tiruchinapalli, M. D. Ehlers, J. D. Keene, Activity-dependent expression of RNA binding protein HuD and its association with mRNAs in neurons. RNA Biol 5, 157–168 (2008).

28. Y. S. Lee et al., AU-rich element-binding protein negatively regulates CCAAT enhancer-binding protein mRNA stability during long-term synaptic plasticity in Aplysia. Proc Natl Acad Sci U S A 109, 15520–15525 (2012).

29. E. Uribe-Querol, C. Rosales, Phagocytosis: Our Current Understanding of a Universal Biological Process. Front Immunol 11, 1066 (2020).

30. Y. Zhu et al., Migratory Neural Crest Cells Phagocytose Dead Cells in the Developing Nervous System. Cell 179, 74–89 e10 (2019).

31. R. C. Paolicelli et al., Synaptic pruning by microglia is necessary for normal brain development. Science 333, 1456–1458 (2011).

32. J. M. Dundee, M. Puigdellivol, R. Butler, G. C. Brown, P2Y(6) Receptor-Dependent Microglial Phagocytosis of Synapses during Development Regulates Synapse Density and Memory. J Neurosci 43, 8090–8103 (2023).

33. L. Weinhard et al., Microglia remodel synapses by presynaptic trogocytosis and spine head filopodia induction. Nat Commun 9, 1228 (2018).

34. I. Diaz-Aparicio et al., Microglia Actively Remodel Adult Hippocampal Neurogenesis through the Phagocytosis Secretome. J Neurosci 40, 1453–1482 (2020).

35. H. Keren-Shaul et al., A Unique Microglia Type Associated with Restricting Development of Alzheimer’s Disease. Cell 169, 1276–1290 e1217 (2017).

36. A. Basu et al., let-7 miRNA controls CED-7 homotypic adhesion and EFF-1-mediated axonal self-fusion to restore touch sensation following injury. Proc Natl Acad Sci U S A 114, E10206–E10215 (2017).

37. C. J. Smith et al., Time-lapse imaging and cell-specific expression profiling reveal dynamic branching and molecular determinants of a multi-dendritic nociceptor in C. elegans. Dev Biol 345, 18–33 (2010).

38. M. Hammarlund, E. M. Jorgensen, M. J. Bastiani, Axons break in animals lacking beta-spectrin. J Cell Biol 176, 269–275 (2007).

39. L. A. McCord et al., Conformational states and recognition of amyloidogenic peptides of human insulin-degrading enzyme. Proc Natl Acad Sci U S A 110, 13827–13832 (2013).

40. J. M. Kinchen et al., Two pathways converge at CED-10 to mediate actin rearrangement and corpse removal in C. elegans. Nature 434, 93–99 (2005).

41. J. Mapes et al., CED-1, CED-7, and TTR-52 regulate surface phosphatidylserine expression on apoptotic and phagocytic cells. Curr Biol 22, 1267–1275 (2012).

42. H. Chiu et al., Engulfing cells promote neuronal regeneration and remove neuronal debris through distinct biochemical functions of CED-1. Nat Commun 9, 4842 (2018).

43. Z. Li et al., Necrotic Cells Actively Attract Phagocytes through the Collaborative Action of Two Distinct PS-Exposure Mechanisms. PLoS Genet 11, e1005285 (2015).

44. N. Hisamoto et al., Phosphatidylserine exposure mediated by ABC transporter activates the integrin signaling pathway promoting axon regeneration. Nat Commun 9, 3099 (2018).

45. H. K. Brar et al., Dendrite regeneration in C. elegans is controlled by the RAC GTPase CED-10 and the RhoGEF TIAM-1. PLoS Genet 18, e1010127 (2022).

46. E. S. Suvorova et al., Discovery of a splicing regulator required for cell cycle progression. PLoS Genet 9, e1003305 (2013).

47. B. M. Ben-Oz et al., A dual role of RBM42 in modulating splicing and translation of CDKN1A/p21 during DNA damage response. Nat Commun 14, 7628 (2023).

48. C. E. Richardson et al., PTRN-1, a microtubule minus end-binding CAMSAP homolog, promotes microtubule function in Caenorhabditis elegans neurons. Elife 3, e01498 (2014).

49. L. E et al., An Antimicrobial Peptide and Its Neuronal Receptor Regulate Dendrite Degeneration in Aging and Infection. Neuron 97, 125–138 e125 (2018).

50. W. C. Duckworth, R. G. Bennett, F. G. Hamel, Insulin acts intracellularly on proteasomes through insulin-degrading enzyme. Biochem Biophys Res Commun 244, 390–394 (1998).

51. R. B. Darnell, RNA protein interaction in neurons. Annu Rev Neurosci 36, 243–270 (2013).

52. C. M. Loya, D. Van Vactor, T. A. Fulga, Understanding neuronal connectivity through the post-transcriptional toolkit. Genes Dev 24, 625–635 (2010).

53. A. S. Hafner, P. G. Donlin-Asp, B. Leitch, E. Herzog, E. M. Schuman, Local protein synthesis is a ubiquitous feature of neuronal pre- and postsynaptic compartments. Science 364, (2019).

