## supplemental data for "Injury-induced nuclear export of RNA-binding proteins drives mRNA stabilization and translation to promote dendrite regeneration"

**The file includes:**

Materials and Methods

References

Figures S1 to S7

Tables S1 to S3

Materials and Methods

Lead Contact and Materials Availability

*C. elegans* strains and maintenance

*C. elegans* were cultured on nematode growth medium (NGM) seeded with *E. coli* OP50 at 20°C following standard protocols (*1*). Bristol N2 strain was used as the wild-type. Transgenic lines were obtained by injecting plasmid DNA mixes into the gonads of hermaphrodites. Integrated transgenic strains were generated using ultraviolet trimethylpsoralen (UV/TMP) treatment and subsequently backcrossed with N2 for at least three times prior to experimental use. A complete list of strains used in this study is provided in Table S1.

Genomic DNA extraction

Genomic DNA for whole-genome sequencing was extracted using the Puregene Cell & Tissue Kit (Qiagen, 158388). Approximately 500 mL of mixed-stage *C. elegans* cultures were collected into 15 mL tubes and washed three times with M9 buffer. Worms were then incubated in fresh M9 buffer on a rotator at room temperature for 2 hours to clear intestinal bacteria, followed by freezing at −80°C overnight. After thawing, 3 mL of cell lysis solution and 15 µL of proteinase K (20 mg/mL) were added, and the samples were incubated at 55°C overnight. Subsequently, 15 µL of RNase A solution was added, and the mixture was rotated at 37°C for 3 hours. Next, 1 mL of protein precipitation solution was added into the tube on ice, the mixture was vortexed vigorously, and centrifuged for 10 minutes at 2000 g. The supernatant was transferred to a new 15 mL tube and mixed with 3 mL isopropanol. Then centrifuge for 3 minutes at 2000 g and discard the supernatant. The resulting pellet was washed with 3 mL of 70% ethanol, centrifuged, air-dried, and dissolved in 150 µL of ultrapure distilled water. Purified genomic DNA samples were submitted to BGI Genomics for whole-genome sequencing.

DNA constructs and generation of transgenes

All constructs used in this study are listed in Table S2. Plasmids used for microinjection were generated by Gateway Cloning technology. DNA fragments of genes or related promoters were amplified by polymerase chain reaction (PCR) and inserted into the pCR8 entry vector (Invitrogen) or destination vector via Gibson assembly. The expression plasmids were obtained by LR recombination reaction (Invitrogen) between entry and destination vectors. Transgenic animals were generated following standard microinjection procedures (*2*). In general, plasmid DNAs of interest were used at 10-100 ng/mL with a co-injection marker, including *Punc-122::RFP::unc-54 3’-UTR* (50 ng/μl), *Pttx-3p::RFP::unc-54 3’-UTR* (50 ng/μl) or *Pf53f4.13::GFP::unc-54 3’-UTR* (20 ng/μl). Rescue strains or RNAi knockdown strains were driven by endogenous promotor (*EP*), PVD-specific promotor (*ser-2(3)*) or hypodermis-specific promotor (*col-19*), respectively.

Genetic screen

The strain NYL3271 was utilized for the dendrite regeneration genetic screen. This strain carries the *unc-70(syb2578syb2747)* (Sunybiotech) and *yadIs171(PVD::FLP-NLS;PVD::GFP)*. In *unc-70(syb2578syb2747)*, one FRT site was inserted into the genomic DNA immediately upstream of the *unc-70* start codon, and the second FRT site was inserted within the third intron (corresponding to the second intron of the *unc-70* isoform c) (Fig. S2A). In *yadIs171(PVD::FLP-NLS;PVD::GFP)*, FLP-NLS and GFP were specifically expressed in PVD neurons driven by *ser-2(3)* promoter. After mutagenesis, we screened phenotypes of defects in dendrite regeneration in the F2 generation from approximate 400 independent F1 lines and identified five mutants showing impaired dendrite regeneration (Fig. 1C).

UV laser dendrotomy and fluorescence microscopy

Day 1 adult animals were used to induce injury on the primary dendrites of PVD neurons (dendrotomy). Animals were anesthetized by using 5 mM levamisole dissolved in M9 buffer and positioned with left side up on a 3% agarose pad. The primary dendrite of each PVD neuron was severed 100–150 µm anterior to the soma using a Zeiss fluorescence microscope equipped with a 435- or 527-nm MicroPoint Laser System (laser dye cells, MP-27-435-DYE). Following dendrotomy, animals were transferred to nematode growth medium (NGM) plates for recovery and subsequently subjected to fluorescence imaging and quantification of dendritic fusion rates.

Confocal images of PVD neurons were acquired using a Zeiss LSM700 confocal microscope. Animals were anesthetized in 5 mM levamisole dissolved in M9 buffer. Detector gain and laser power were optimized to minimize pixel saturation and maximize the dynamic detection range. Images presented in the figures represent maximum-intensity projections of z-stack acquisitions obtained under a 40× objective.

Real-time quantitative reverse-transcription PCR (RT-qPCR)

To quantify *ced-7* mRNA levels, total RNA was extracted from wild-type N2 (Control), *idr-1(lf)*, *rbm-42(lf)*, and *rbm-42(gf)* animals cultured on NGM plates. RNA was isolated using TRI Reagent and purified with the Direct-zol RNA MiniPrep Kit (Zymo Research, R11330). Reverse transcription was performed with the iScript Reverse Transcription Supermix for RT-qPCR (Bio-Rad, 1708840), followed by quantitative PCR using iTaq Universal SYBR Green Supermix (Bio-Rad, 1725120) on a StepOnePlus™ Real-Time PCR System (Applied Biosystems). *Actin* served as the endogenous control. Three independent biological replicates were analyzed. Relative mRNA levels were quantified using StepOnePlus™ software V2.3 (Applied Biosystems) based on the comparative ∆CT method. Primers used for amplifying *ced-7* and *actin* were listed in Table S3.

Quantification of dendrite regeneration

To evaluate dendrite regeneration, we quantified the dendritic fusion rate, which serves as the principal indicator of recovery in PVD neurons (*3*). Our results confirmed that dendritic fusion corresponds to both morphological and functional reconnection of severed dendrites (Fig. 1A, Fig. S1A and 1B).

For quantification, Day 1 adult animals were subjected to laser dendrotomy as described above. At defined time points post-injury (typically 24–96 hours), animals were transferred on 3% agarose pads in M9 buffer, and PVD neurons were visualized under the compound microscope. The dendritic fusion rate was calculated as the percentage of animals exhibiting successful fusion among the total number of animals examined. Each experiment included at least three independent replicates.

PVD neuron-specific RNAi

To knock down gene expression specifically in PVD neurons, DNA fragments corresponding to the target genes (*rbm-42* and *ced-7*) were PCR-amplified and cloned in both forward and reverse orientations into the pCR8 entry vector (Invitrogen) separately. These fragments were subsequently recombined into destination vectors containing a PVD neuron–specific promoter using Gateway cloning. The resulting plasmids with both directions were mixed and microinjected into adult hermaphrodites to generate transgenic animals expressing double-stranded RNA under PVD-specific control. Day 1 adult transgenic animals were then subjected to laser dendrotomy, and dendritic fusion rates were quantified at the indicated time points. Primers used for amplifying fragments of *rbm-42* and *ced-7* were listed in Table S3.

Statistic

Data were analyzed using paired Student’s *t* test, chi-square test, one-way analysis of variance (ANOVA) and two-way ANOVA followed by Tukey’s post hoc test in GraphPad Prism 10. A *P* value < 0.05 was considered statistically significant. Data are presented as means ± standard deviation (SD). For all dendrotomy experiments, *n* in the figure legends indicates the number of animals per condition in each experiment, and statistics were derived from three independent biological experiments.

References

1.	S. Brenner, The genetics of Caenorhabditis elegans. *Genetics* **77**, 71-94 (1974).

2.	C. C. Mello, J. M. Kramer, D. Stinchcomb, V. Ambros, Efficient gene transfer in C.elegans: extrachromosomal maintenance and integration of transforming sequences. *EMBO J* **10**, 3959-3970 (1991).

3.	M. Oren-Suissa, T. Gattegno, V. Kravtsov, B. Podbilewicz, Extrinsic Repair of Injured Dendrites as a Paradigm for Regeneration by Fusion in Caenorhabditis elegans. *Genetics* **206**, 215-230 (2017).

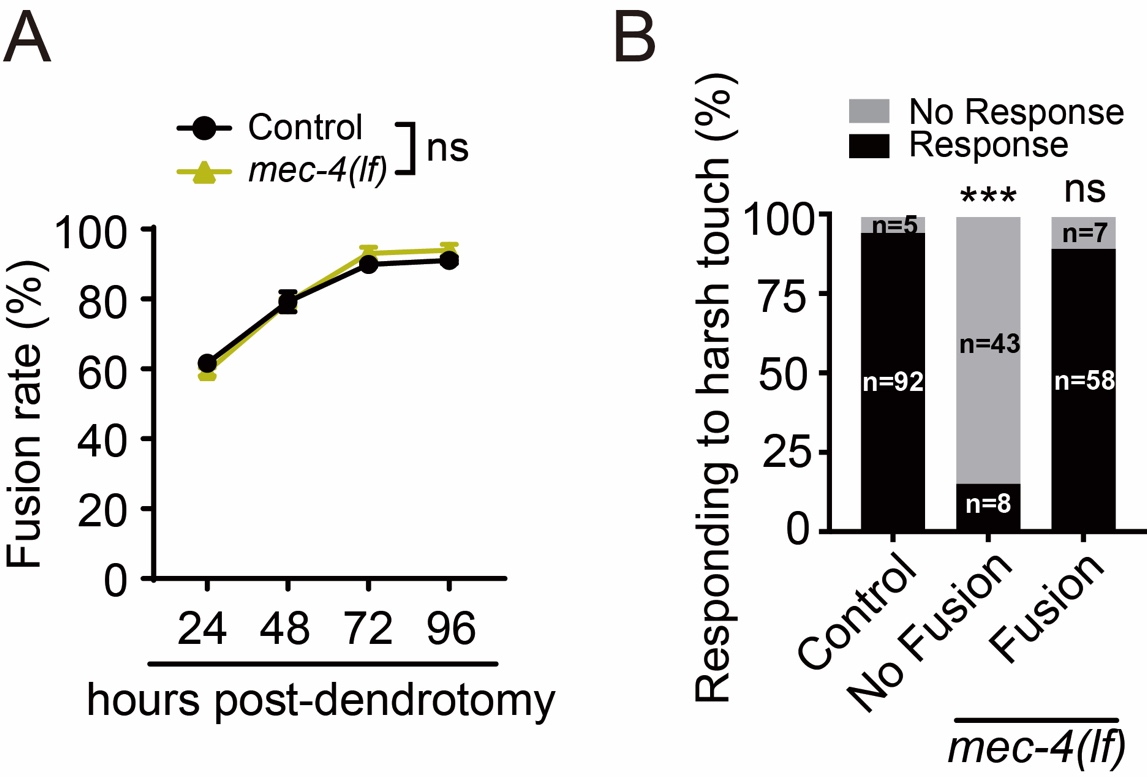

Figure S1. Dendritic fusion is functional recovery.

(**A**) Quantification data show that *mec-4(lf)* does not change the dendritic fusion rate after dendrotomy. Dendrotomy was performed on 1-day-old adult animals. *n*≥32. Data represent mean ± standard deviation (SD) from three independent experiments. ns, nonsignificant. Statistical significance was determined by two-way ANOVA followed by Tukey’s post hoc test.

(**B**) Comparison of harsh touch responses between control and injured animals. Percentage of animals responding to harsh touch was compared among control animals and *mec-4(lf)* mutants exhibiting either fusion or non-fusion phenotypes following injury. ****P* ≤ 0.001; ns, nonsignificant. Statistical significance was determined by chi-square test.

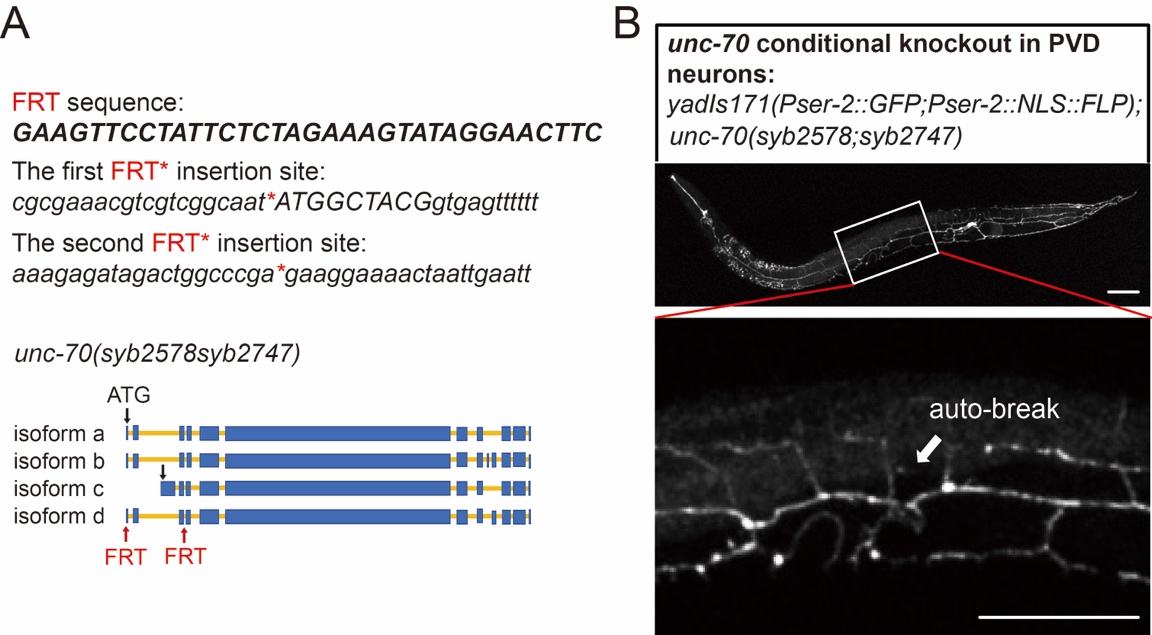

Figure S2. Generation of the strain for forward genetic screen.

(A) *unc-70* FRT knock-in. The FRT sequence is shown in the upper panel, and the corresponding insertion sites are indicated in the lower panel. Two FRT fragments were inserted right before *unc-70* start codon and in the third intron (the second intron of isoform b), respectively.

(B) Representative images of an animal with *unc-70* conditionally knocked out in PVD neurons. The region showing an auto-break in the primary dendrite is enlarged and displayed below. White arrows indicate axonal auto-breaks. Scale bar, 100 µm.

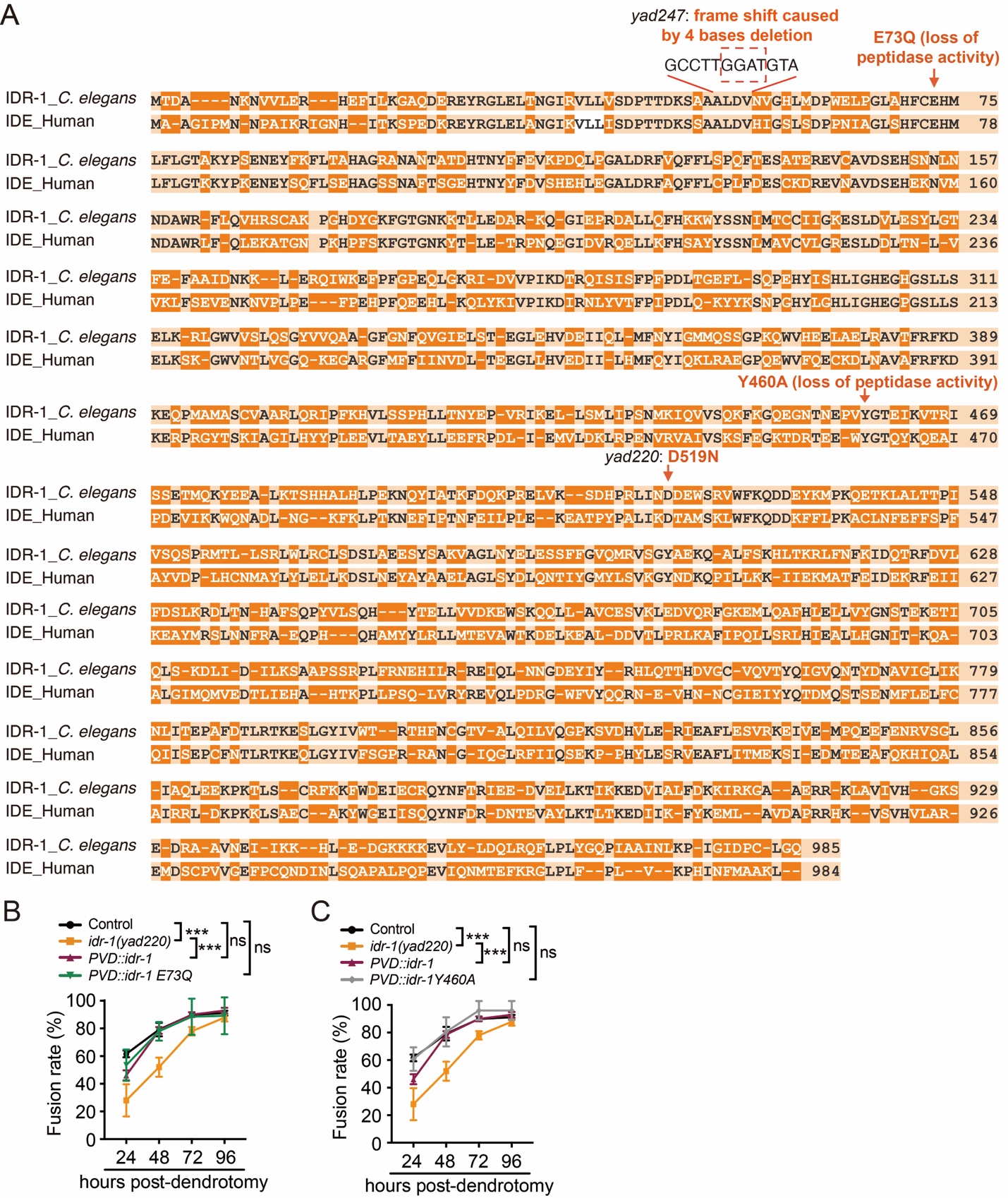

**Figure S3.** **Sequence alignment of *C. elegans* IDR-1 and mammalian IDE.**

(A) Sequence alignment of *C. elegans* IDR-1 and human IDE. The alleles *yad220* and *yad247* are labeled. Residues critical for peptidase activity, E73Q and Y460A, are indicated.

(B and C) Quantification of dendritic fusion rate in Control, *idr-1(lf)* and rescue strains expressing *idr-1*, *idr-1E73Q* (B) or *idr-1Y460A* (C) specifically in PVD neurons. *n*≥30. Data represent mean ± SD from three independent experiments. ****P* ≤ 0.001; ns, nonsignificant. Statistical significance was determined by two-way ANOVA followed by Tukey’s post hoc test.

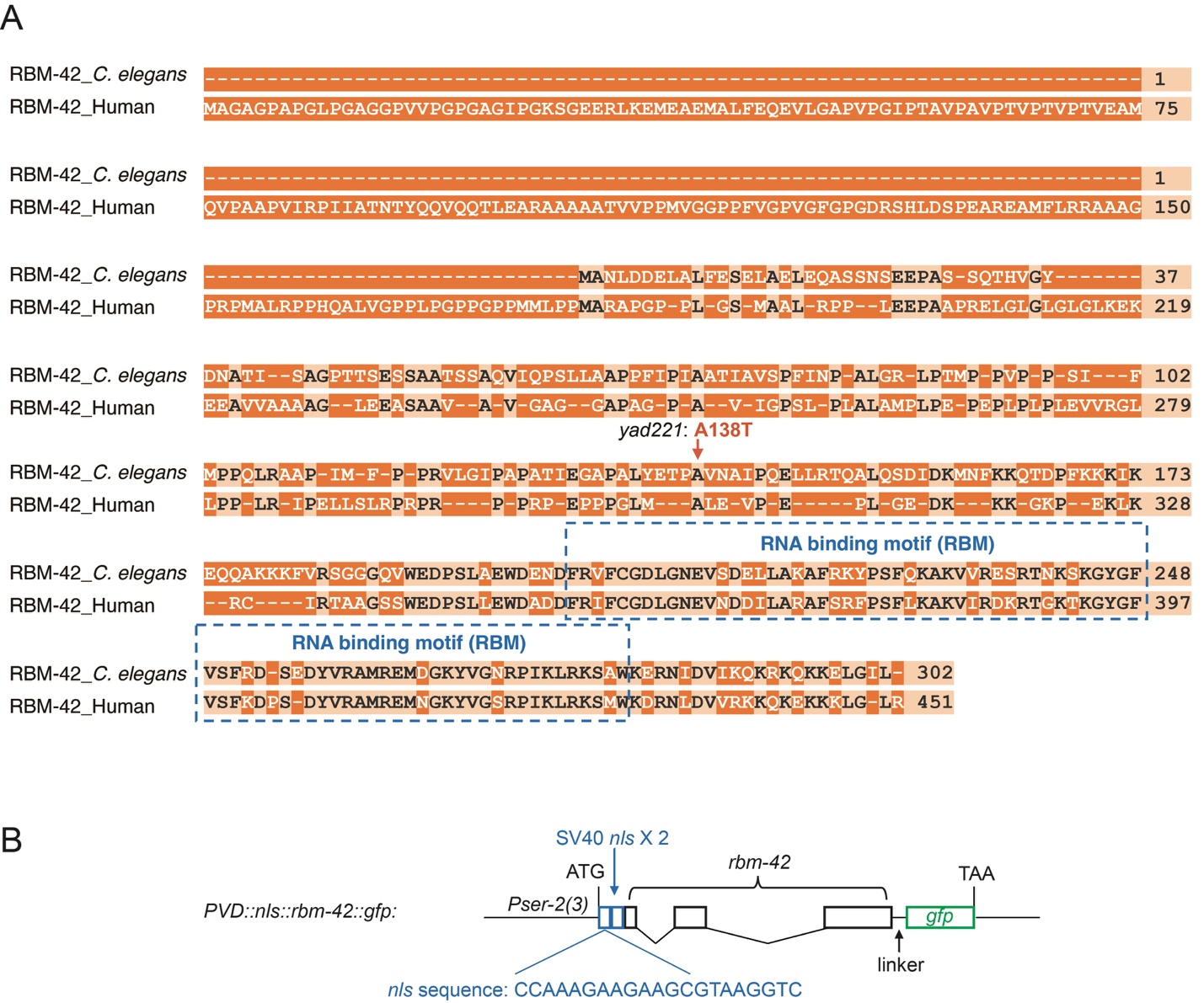

**Figure S4.** **Sequence alignment of *C. elegans* RBM-42 and mammalian RBM42.**

(A) Sequence alignment of *C. elegans* RBM-42 and human RBM42. The allele *yad221* and RNA-binding motif are labeled.

(B) Schematic illustration of the *PVD::nls::rbm-42::gfp* construct, showing the arrangement of *ser-2(3)* promotor, two tandem SV40 nuclear localization sequences (NLS), the *rbm-42* coding region, a linker, and the *gfp* sequence.

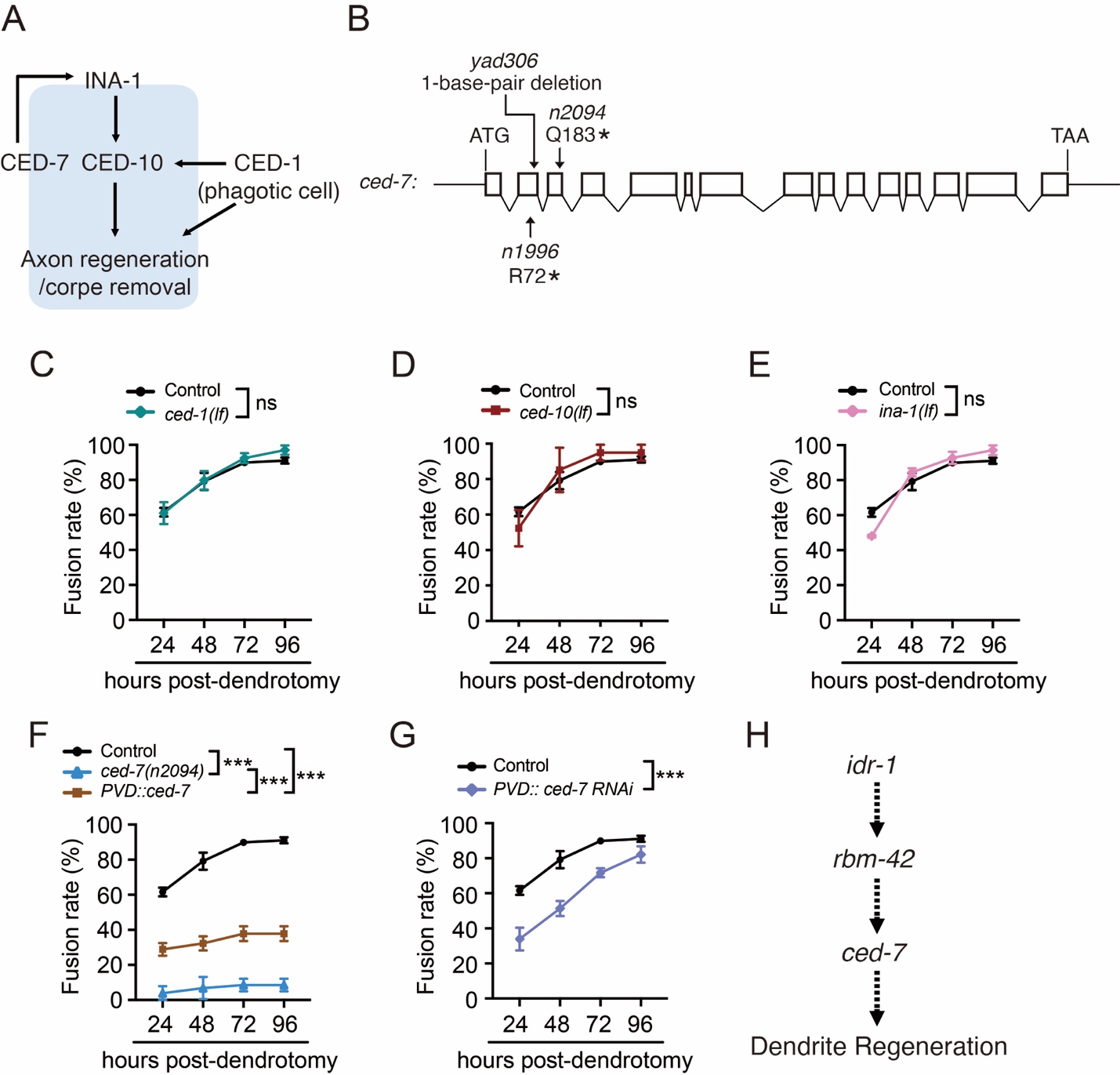

**Figure S5. *ced-7*, but not other phagocytosis-related genes, is required for dendrite regeneration.**

(**A**) A diagram showing the functional relationships of *ced-1*, *ced-7*, *ced-10*, and *ina-1* in axonal regeneration and cell corpse clearance.

(**B**) A schematic illustrating the nature of *ced-7* mutations. *n1996* and *n2094* introduce premature stop codons, whereas *yad306* causes a frameshift mutation and premature stop codon.

(**C**-**E**) Comparison of dendritic fusion rate between control and *ced-1(lf)* (C), *ced-10(lf)* (D) and *ina-1(lf)* (E) animals at 24, 48, 72, and 96 hours post-dendrotomy. *n*≥24.

(**F**) Expression of *ced-7* in PVD neurons rescues dendrite regeneration defect in *ced-7(n2094)* animals. Quantification of dendritic fusion rate in control, *ced-7(n2094)* and rescue strain. *n*≥24.

(**G**) Knockdown of *ced-7* specifically in PVD neurons suppresses dendrite regeneration. Quantification data of dendritic fusion rate in control and *PVD::ced-7 RNAi* animals. *n*≥3**2.**

(**H**) Schematic illustrating the genetic relationships among *idr-1*, *rbm-42* and *ced-7* in dendrite regeneration.

Data represent mean ± SD from three independent experiments. ****P* ≤ 0.001; ns, nonsignificant. Statistical significance was determined by Two-way ANOVA followed by Tukey’s post hoc test.

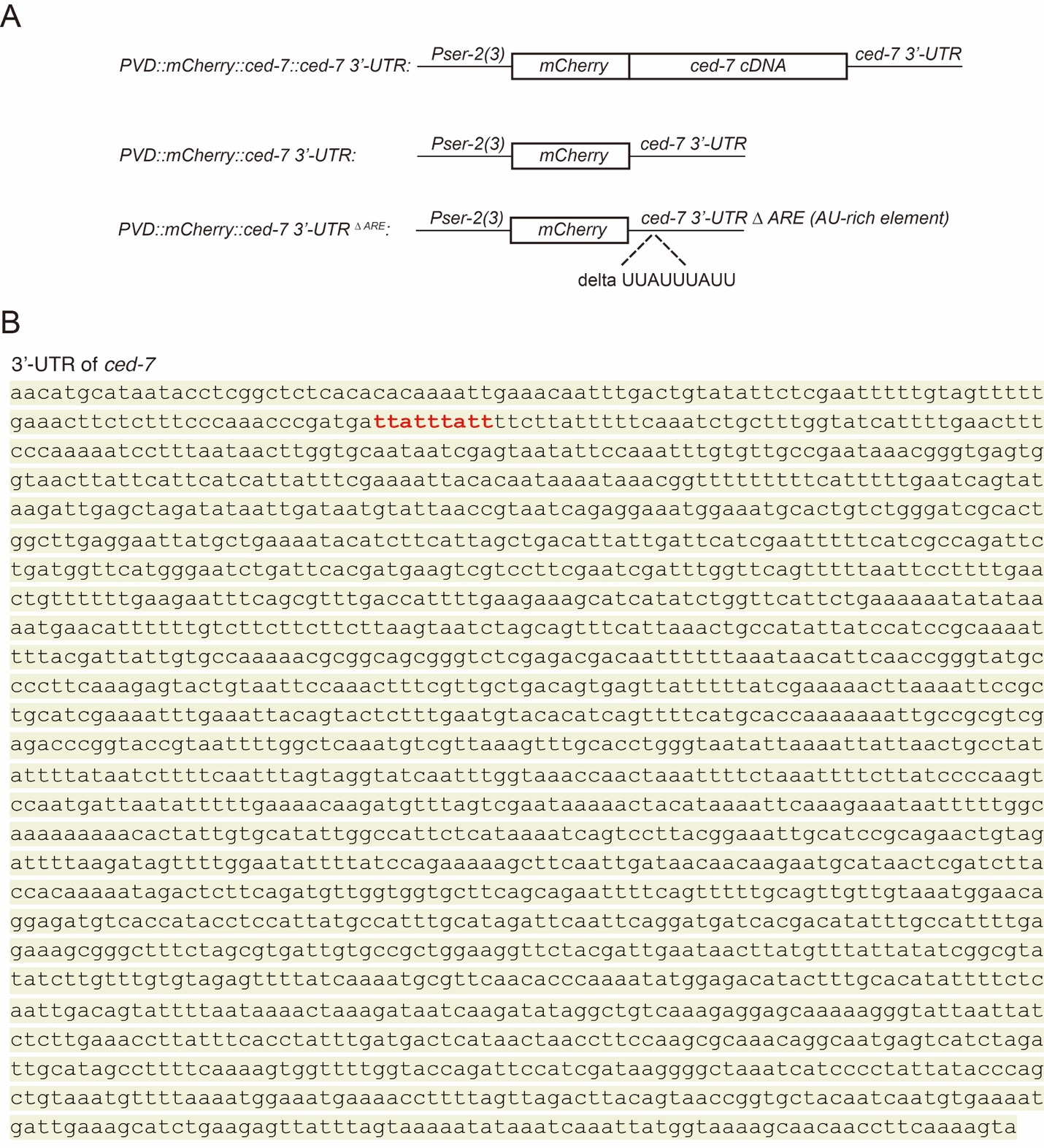

**Figure S6. Diagram of expression constructs of mCherry fused with 3’-UTR of *ced-7* used for local translation experiments.**

(**A**) Diagram of expression constructs used in Fig. 6C, 6H and 6M.

(**B**) The adenine/uridine-rich element (ARE) in 3’-UTR of *ced-7* is highlighted in red.

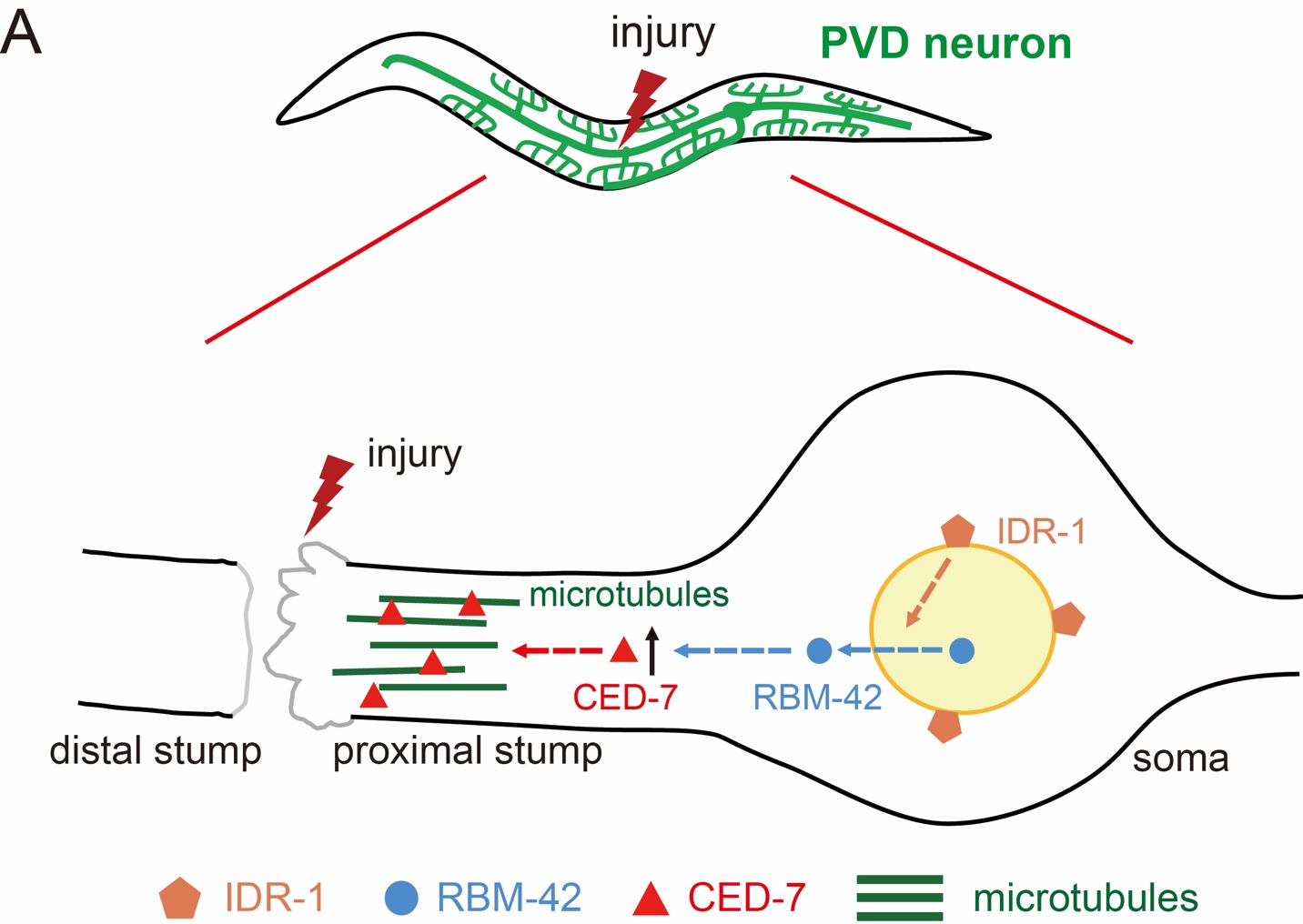

**Figure S7. A model for *idr-1-rbm-42-ced-7* signal cascade in mediating dendrite regeneration.**

(**A**) Following injury, activation of IDR-1 triggers the nuclear export of RBM-42, which in turn enhances CED-7 translation. Upregulation of CED-7 facilitates microtubule assembly and promotes dendrite regeneration.

| Table 1. *C. elegans* strains made in this study | | |
| --- | --- | --- |
| Strain name | Genotype | Related Figures |
| NYL1116 | *mec-4(u253);wdIs51[F49H12.5::GFP+unc-119(+)]* | Fig. S1A |
| PHX2747 | *unc-70(syb2747syb2758)* | Fig. S2A |
| NYL3271 | *yadIs171[Pser-2(3)::GFP; Pser-2(3)::NLS::FLP; Pf53f4.13::mCherry]; unc-70(syb2747syb2758)* | Fig. 1C |
| NYL3669 | *idr-1(yad220);wdIs51[F49H12.5::GFP+unc-119(+)]* | Fig. 1E-1H, Fig. S3B and S3C |
| NYL3683 | *idr-1(yad220);wdIs51[F49H12.5::GFP+unc-119(+)]; yadEx1793[Pidr-1::idr-1; Punc-122::RFP]* | Fig. 1E |
| NYL3876 | *idr-1(yad247);wdIs51[F49H12.5::GFP+unc-119(+)]* | Fig. 1E, Fig. 4A, 4B and Fig. 4E |
| NYL4004 | *idr-1(yad220);wdIs51[F49H12.5::GFP+unc-119(+)]; yadEx1995[Pser-2(3)::idr-1; Punc-122::RFP]* | Fig. 1F and 1G |
| NYL4006 | *idr-1(yad220);wdIs51[F49H12.5::GFP+unc-119(+)]; yadEx1997[Pcol-19::idr-1; Punc-122::RFP]* | Fig. 1F |
| NYL3878 | *idr-1(yad220);wdIs51[F49H12.5::GFP+unc-119(+)]; yadEx1966[Pser-2(3)::IDE(human); Punc-122::RFP]* | Fig. 1G |
| NYL5024 | *idr-1(yad220);wdIs51[F49H12.5::GFP+unc-119(+)]; yadEx2506[Pser-2(3)::idr-1 E73Q; Punc-122::RFP]* | Fig. S3B |
| NYL5025 | *idr-1(yad220);wdIs51[F49H12.5::GFP+unc-119(+)]; yadEx2507[Pser-2(3)::idr-1 Y460A; Punc-122::RFP]* | Fig. S3C |
| NYL3673 | *rbm-42(yad221);wdIs51[F49H12.5::GFP+unc-119(+)]* | Fig. 2B-2F, Fig. 4A-4D and Fig. 6A and 6B |
| NYL3681 | *rbm-42(yad221);wdIs51[F49H12.5::GFP+unc-119(+)]; yadEx1791[Prbm-42::rbm-42; Punc-122::RFP]* | Fig. 2B, Fig. 4E and 4F |
| NYL5027 | *wdIs51[F49H12.5::GFP+unc-119(+)];yadEx2468[Pser-2(3)::rbm-42 RNAi; Punc-122::RFP]* | Fig. 2C |
| NYL3797 | *rbm-42(yad221);wdIs51[F49H12.5::GFP+unc-119(+)]; yadEx1843[Pser-2(3)::rbm-42; Punc-122::RFP]* | Fig. 2D and Fig. 6A |
| NYL4103 | *rbm-42(yad221);wdIs51[F49H12.5::GFP+unc-119(+)]; yadEx2157[Pcol-19::rbm-42; Punc-122::RFP]* | Fig. 2D |
| NYL4008 | *rbm-42(yad221);wdIs51[F49H12.5::GFP+unc-119(+)]; yadEx1999[Pser-2(3)::RBM42; Punc-122::RFP]* | Fig. 2E |
| NYL4871 | *ced-7(n1996);wdIs51[F49H12.5::GFP+unc-119(+)]* | Fig. 3A-3C, Fig. 4C, 4D and 4F |
| NYL4820 | *ced-7(n2094);wdIs51[F49H12.5::GFP+unc-119(+)]* | Fig. 3A and Fig. S5F |
| NYL4803 | *ced-1(e1735);wdIs51[F49H12.5::GFP+unc-119(+)]* | Fig. S5C |
| NYL4230 | *ced-10(n3246);wdIs51[F49H12.5::GFP+unc-119(+)]* | Fig. S5D |
| NYL4809 | *ina-1(gm144);wdIs51[F49H12.5::GFP+unc-119(+)]* | Fig. S5E |
| NYL4985 | *ced-7(n1996);wdIs51[F49H12.5::GFP+unc-119(+)]; yadEx2471[Pser-2(3)::ced-7::ced-7 3’-UTR; Punc-122::RFP]* | Fig. 3B |
| NYL4998 | *ced-7(n1996);wdIs51[F49H12.5::GFP+unc-119(+)]; yadEx2475[Pcol-19::ced-7::ced-7 3’-UTR; Punc-122::RFP]* | Fig. 3B |
| NYL5071 | *ced-7(n2094);wdIs51[F49H12.5::GFP+unc-119(+)]; yadEx2479[Pser-2(3)::ced-7::ced-7 3’-UTR; Punc-122::RFP]* | Fig. S5F |
| NYL4922 | *wdIs51[F49H12.5::GFP+unc-119(+)];yadEx2483[Pser-2(3)::ced-7 RNAi; Punc-122::RFP]* | Fig. S5G |
| NYL3985 | *idr-1(yad220); rbm-42(yad221); wdIs51[F49H12.5::GFP+unc-119(+)]* | Fig. 4A and 4B |
| NYL5110 | *rbm-42(yad221)ced-7(yad306); wdIs51[F49H12.5::GFP+unc-119(+)]* | Fig. 4C and 4D |
| NYL4131 | *wdIs51[F49H12.5::GFP+unc-119(+)];yadEx1791[Prbm-42::rbm-42; Punc-122::RFP]* | Fig. 4E and 4F |
| NYL4131 | *idr-1(yad247);wdIs51[F49H12.5::GFP+unc-119(+)]; yadEx1791[Prbm-42::rbm-42; Punc-122::RFP]* | Fig. 4E |
| NYL5026 | *ced-7(n1996);wdIs51[F49H12.5::GFP+unc-119(+)]; yadEx1791[Prbm-42::rbm-42; Punc-122::RFP]* | Fig. 4F |
| NYL5087 | *wdIs51[F49H12.5::GFP+unc-119(+)];yadEx2488[Pser-2(3)::mCherry;ser-2(3)::rbm-42::GFP; Pf53f4.13::GFP]* | Fig. 5A |
| NYL5027 | *idr-1(220); wdIs51[F49H12.5::GFP+unc-119(+)]; yadEx2488[Pser-2(3)::mCherry;ser-2(3)::rbm-42::GFP; Pf53f4.13::GFP]* | Fig. 5A |
| NYL5081 | *rbm-42(yad221);wdIs51[F49H12.5::GFP+unc-119(+)]; yadEx2492[Pser-2(3)::rbm-42::GFP; Pf53f4.13::GFP]* | Fig. 5E |
| NYL5085 | *rbm-42(yad221);wdIs51[F49H12.5::GFP+unc-119(+)]; yadEx2496[Pser-2(3)::nls::rbm-42::GFP; Pf53f4.13::GFP]* | Fig. 5E |
| NYL3900 | *rbm-42(yad221);wdIs51[F49H12.5::GFP+unc-119(+)]; yadEx2492[Pser-2(3)::rbm-42 ∆RBM; Punc-122::RFP]* | Fig. 6A |
| NYL4625 | *rbm-42(yadIs296) [Prbm-42::rbm-42(gf)+ Pf53f4.13::GFP]; wdIs51[F49H12.5::GFP+unc-119(+)]* | Fig. 6B |
| NYL5033 | *wdIs51[F49H12.5::GFP+unc-119(+)]; yadEx2498[Pser-2(3)::mCherry::ced-7::ced-7 3’-UTR; Pf53f4.13::GFP]* | Fig. 6C |
| NYL5030 | *rbm-42(yad221); wdIs51[F49H12.5::GFP+unc-119(+)]; yadEx2498[Pser-2(3)::mCherry::ced-7::ced-7 3’-UTR; Pf53f4.13::GFP]* | Fig. 6C |
| NYL5007 | *wdIs51[F49H12.5::GFP+unc-119(+)];yadEx2501[Pser-2(3)::mCherry::ced-7 3’-UTR; Pf53f4.13::GFP]* | Fig. 6H |
| NYL5002 | *rbm-42(yad221);wdIs51[F49H12.5::GFP+unc-119(+)]; yadEx2501[Pser-2(3)::mCherry::ced-7 3’-UTR; Pf53f4.13::GFP]* | Fig. 6H |
| NYL5028 | *wdIs51[F49H12.5::GFP+unc-119(+)]; yadEx2508[Pser-2(3)::mCherry::ced-7 3’-UTR ∆ARE; Pf53f4.13::GFP]* | Fig. 6M |
| NYL5029 | *rbm-42(yad221);wdIs51[F49H12.5::GFP+unc-119(+)]; yadEx2508[Pser-2(3)::mCherry::ced-7 3’-UTR ∆ARE; Pf53f4.13::GFP]* | Fig. 6M |
| NYL4826 | *yadIs261[Pser-2(3)::mCherry];yadEx2505[Pser-2(3)::EMTB::GFP; Pttx-3::RFP]* | Fig. 7A and 7B |
| NYL4835 | *idr-1(yad220);yadIs261[Pser-2(3)::mCherry]; yadEx2505[Pser-2(3)::EMTB::GFP; Pttx-3::RFP]* | Fig. 7B |
| NYL4836 | *rbm-42(yad221);yadIs261[Pser-2(3)::mCherry]; yadEx2505[Pser-2(3)::EMTB::GFP; Pttx-3::RFP]* | Fig. 7B |
| NYL4834 | *ced-7(n1996);yadIs261[Pser-2(3)::mCherry]; yadEx2505[Pser-2(3)::EMTB::GFP; Pttx-3::RFP]* | Fig. 7B |

Table S2. DNA constructs used in this study

| **Recombinant DNA** | **Source** | **Identifier** |
| --- | --- | --- |
| *Pser-2(3)::idr-1::unc-54 3’-UTR* | This paper | PNYL1835 |
| *Pcol-19::idr-1::unc-54 3’-UTR* | This paper | PNYL1836 |
| *Pser-2(3)::IDE(human)::unc-54 3’-UTR* | This paper | PNYL1581 |
| *Pser-2(3)::idr-1 E73Q::unc-54 3’-UTR* | This paper | PNYL1840 |
| *Pser-2(3)::idr-1 Y460A::unc-54 3’-UTR* | This paper | PNYL1838 |
| *Pser-2(3)::rbm-42 RNAi-1F::unc-54 3’-UTR* | This paper | PNYL1892 |
| *Pser-2(3)::rbm-42 RNAi-1R::unc-54 3’-UTR* | This paper | PNYL1893 |
| *Pser-2(3)::rbm-42 RNAi-2F::unc-54 3’-UTR* | This paper | PNYL1894 |
| *Pser-2(3)::rbm-42 RNAi-2R::unc-54 3’-UTR* | This paper | PNYL1895 |
| *Pser-2(3)::rbm-42::unc-54 3’-UTR* | This paper | PNYL1573 |
| *Pcol-19:: ::rbm-42::unc-54 3’-UTR* | This paper | PNYL1572 |
| *Pser-2(3)::RBM42(human)::unc-54 3’-UTR* | This paper | PNYL1845 |
| *Pser-2(3)::ced-7::ced-7 3’-UTR* | This paper | PNYL1901 |
| *Pcol-19::ced-7::ced-7 3’-UTR* | This paper | PNYL1902 |
| *Pser-2(3)::ced-7 RNAi-F::unc-54 3’-UTR* | This paper | PNYL1905 |
| *Pser-2(3)::ced-7 RNAi-R::unc-54 3’-UTR* | This paper | PNYL1907 |
| *Pser-2(3)::rbm-42::GFP::unc-54 3’-UTR* | This paper | PNYL1922 |
| *Pser-2(3)::mCherry::unc-54 3’-UTR* | This paper | PNYL1924 |
| *Pser-2(3)::nls::rbm-42::GFP::unc-54 3’-UTR* | This paper | PNYL1919 |
| *Pser-2(3)::rbm-42^∆RBM^::unc-54 3’-UTR* | This paper | PNYL1926 |
| *Pser-2(3)::mCherry::ced-7::ced-7 3’-UTR* | This paper | PNYL1910 |
| *Pser-2(3)::mCherry::ced-7 3’-UTR* | This paper | PNYL1900 |
| *Pser-2(3)::mCherry::ced-7 3’-UTR^∆ARE^* | This paper | PNYL1912 |
| *Pser-2(3)::EMTB::GFP::unc-54 3’-UTR* | This paper | PNYL931 |

Table S3. Primers used in this study

| *C.elegans ced-7*  (for RT-qPCR) | Forward: 5’-TCCAAGCGTTAGAGGCCATTC-3’ |
| --- | --- |
|  | Reverse: 5’-CTTGTTCCAGAGTGTTGAGTG-3’ |
| *C.elegans actin*  (for RT-qPCR) | Forward: 5’-TGGGACAGAAAGACTCGTAC-3’ |
|  | Reverse: 5’-TTTCCATGTCGTCCCAGTTG-3’ |
| *C.elegans rbm-42*  (for RNAi-1) | Forward: 5’-TTAGCCGAACTTGAACAAGC-3’ |
|  | Reverse: 5’-GGTGCAGCTCTTAATTGTGG-3’ |
| *C.elegans rbm-42*  (for RNAi-2) | Forward: 5’-AGGTGCTCCAGCATTATATGAAAC-3’ |
|  | Reverse: 5’-CCGTCCATTTCACGCATAGC-3’ |
| *C.elegans ced-7*  (for RNAi) | Forward: 5’-TCTTCTGTGCAATCTGGAGC-3’ |
|  | Reverse: 5’-CAACGACCAAAGCACCAATG-3’ |
